# Attentional brain rhythms during prolonged cognitive activity

**DOI:** 10.1101/2021.05.26.445730

**Authors:** C. Gaillard, C. De Sousa, J. Amengual, C. Loriette, C. Ziane, S. Ben Hadj Hassen, F. Di Bello, S. Ben Hamed

## Abstract

As routine and lower demand cognitive tasks are taken over by automated assistive systems, human operators are increasingly required to sustain cognitive demand over long periods of time. This has been reported to have long term adverse effects on cardiovascular and mental health. However, it remains unclear whether prolonged cognitive activity results in a monotonic decrease in the efficiency of the recruited brain processes, or whether the brain is able to sustain functions over time spans of one hour and more. Here, we show that during working sessions of one hour or more, contrary to the prediction of a monotonic decline, behavioral performance in both humans and non-human primates consistently fluctuates between periods of optimal and suboptimal performance at a very slow rhythm of *circa* 5 cycles per hour. These fluctuations are observed in both high attentional (in non-human primates) and low attentional (in humans) demand conditions. They coincide with fluctuations in pupil diameter, indicating underlying changes in arousal and information-processing load. Accordingly, we show that these rhythmic behavioral fluctuations correlate, at the neurophysiological level, with fluctuations in the informational attention orientation and perception processing capacity of prefrontal neuronal populations. We further identify specific markers of these fluctuations in LFP power, LFP coherence and spike-field coherence, pointing towards long-range rhythmic modulatory inputs to the prefrontal cortex rather than a local prefrontal origin. These results shed light on the resilience of brain mechanisms to sustained effort and have direct implications on how to optimize high cognitive demand working and learning environments.

## Introduction

A six hours long university exam or a routine maneuver at the international space station require highly sustained performance and any drop in cognition can have dramatic outcomes. We are increasingly faced with such situations in which memory and decision-making processes need to be sustained at high performance for long time periods. However, maintaining cognitive processes at an optimal regime comes at a cost. For example, prolonged activity has long term adverse effects on health, ranging from a higher rate of cardiovascular diseases to fatigue, reduced sleep duration and depression(van der Hulst, 2003; Johnson and Lipscomb, 2006; Liu and Tanaka, 2002; Sekine et al., 2006; Shields, 1999; Sokejima and Kagamimori, 1998). In addition, prolonged cognitive activity also results in a decrease in cognitive performance associated with undesirable decision-making, distress and burnout(Lockley et al., 2004; Proctor et al., 1996; Virtanen et al., 2009). In particular, sustained attention over long time periods has been linked to mental fatigue and cognitive exhaustion(Bonnefond et al., 2010; Guo et al., 2016; Warm et al., 2008).

The short term influence of motivational factors such as reward history(Abrahamyan et al., 2016; Urai et al., 2019), or error history(Cowley et al., 2020; Pisupati et al.; Weissman et al., 2006), on decisions and neuronal processes have been extensively studied and modelled. In contrast, the long-term influence of task-independent factors such as fatigue(Marcora et al., 2009), arousal(Aston-Jones and Cohen, 2005; McGinley et al., 2015; Urai et al., 2019), and satiety(Allen et al., 2019), are less understood. In particular, the neuronal mechanisms associated with sustained cognitive performance over prolonged periods of time are to date poorly understood and it remains unclear whether prolonged cognitive activity results in a monotonic decrease in the efficiency of the recruited brain processes, or whether the brain is actually able to sustain functions over time spans of one hour and more. A recent study in non-human primates reports that, during prolonged cognitive tasks, a slow drift in the neuronal activity of the visual and prefrontal cortex is associated with enhanced impulsive decisions(Cowley et al., 2020).

In the following, we performed a cross-species study in which we show that during working sessions of one hour or more, behavioral performance in both humans and non-human primates, consistently fluctuates at a very slow rhythm of the order of 5 cycles per hour. Importantly, we show that these behavioral fluctuations are qualitatively similar in both species (humans and macaques), during the performance of tasks involving different levels of attentional engagement. We characterize the neurophysiological correlates of these ultra-slow behavioral fluctuations by analyzing variations in pupil size in humans and variations in prefrontal perceptual and attention neuronal processes in non-human primates.

Specifically, we trained two macaque monkeys to perform a very long sustained attention task lasting from one to four hours. We simultaneously recorded multi-unit neuronal activity (MUA) and local field potentials (LFP), from multiple recording sites in the frontal eye field (FEF), a prefrontal cortical area at the source of spatial attention control signals(Astrand et al., 2016; Buschman and Miller, 2007a; Ekstrom et al., 2008; Gregoriou et al., 2009a; Ibos et al., 2013). We report large fluctuations in attentional behavioral performance, by up to 10%, at an ultra-slow rhythm of 4 to 7 cycles per hour (i.e. every 9 to 15 minutes). Applying machine learning approaches to decode prefrontal attentional related information(Astrand et al., 2016, 2020; De Sousa et al., 2021; Gaillard et al., 2020a), we show that these fluctuations in behavioral performance coincide with phase-locked rhythmic fluctuations in how well the prefrontal cortex both accurately encodes the perception of incoming visual stimuli and spatial attention orientation information. In addition, we show that high behavioral performance task epochs coincide with the enhanced of multiple neurophysiological attentional markers: enhanced theta (4-6Hz) and beta (18-24Hz) LFP power, enhanced alpha (7-11Hz) LFP coherence, enhanced spike-field coherence in the theta (4-6Hz), alpha (7-11Hz) and gamma (35-50Hz) frequencies. In humans, we further show that these fluctuations in behavioral performance coincide with rhythmic variations in pupil size, pointing towards underlying changes in arousal and information-processing load. Overall, this suggest that the hour-scale variations in cognitive performance arise from long range mechanisms, impacting local neuronal spiking activity through selective changes in spike-field coherence. The net effect is an overall variation in the efficiency of brain attention and perceptual related processes. These results are discussed in the context of perceptual and attentional rhythms and the resilience of brain mechanisms to sustained effort. In addition to shedding light on how the brain functions under prolonged cognitive activity, this work has direct implications on how to optimize high cognitive demand working and learning environments to reconcile short term effort with both long-term performance and mental health.

## Results

In order to describe modulations of attentional behavioral performance, and their prefrontal neuronal correlates, we had monkeys and humans perform a cognitive task for very long durations while respectively either recoding prefrontal neuronal activity or pupil size. Specifically, we had two monkeys perform a demanding attentional task for one to four hours, depending on the motivation of the monkeys. Monkeys were required to perform a cued target-detection task (Fig. 1a) while we recorded the MUA and LFP activity bilaterally the frontal eye fields (FEF), using two 24-contacts recording probes (Fig. 1b). Target luminosity was adjusted so as to enhance the attentional relevance of the cue, and distractors were introduced in order to force the monkeys to only respond to the target. Fig. 1c shows the behavioral performance of both monkeys during this cued target-detection task. Median task duration was 3.94 hours for monkey M1 (s.e. = 0.66) and 2.99 hours for monkey M2 (s.e. = 0.20). Overall, behavioral performance did not vary between the beginning (defined as the initial 20 minutes of the task) and the end of the behavioral session (defined as the last 20 minutes of the task, Wilcoxon non-parametric test, p=0.40), which indicates that we did not find a trend of decreasing performance along the session. Human subjects (35 subjects in all), had to detect a shivering Gabor in the midst of slowly drifting Gabors of variable orientations. Task duration in human subjects was fixed to 75 min. As in macaques, there was no significant changes in median reaction times when comparing the first versus the last 20 minutes of the task (Wilcoxon non-parametric test, p=0.95).

**Figure 1:**
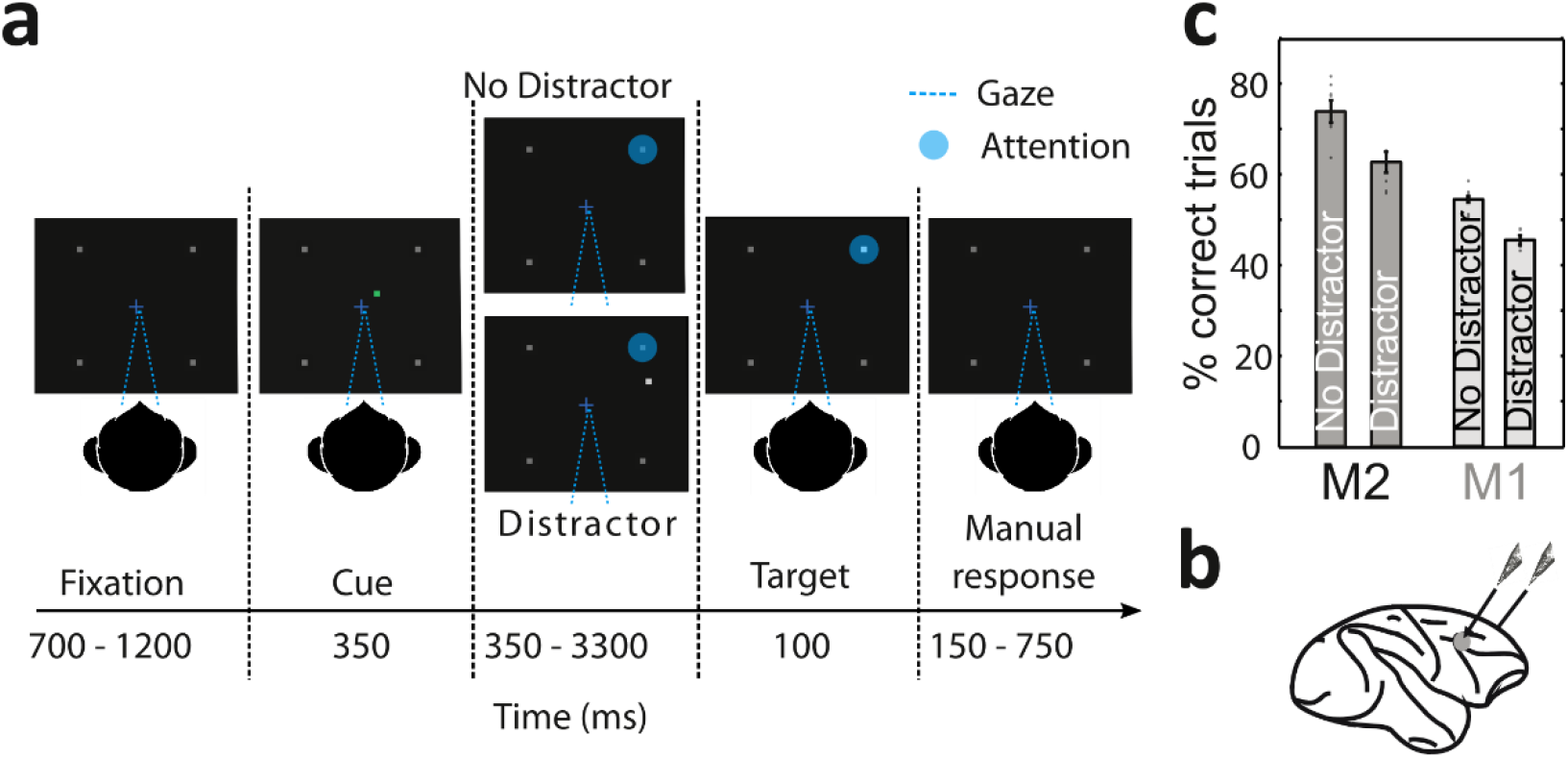
Task design and behavioral performance. (A) 100% validity cued target detection task with temporal distractors. Monkeys had to hold a bar and fixate a central cross on the screen for a trial to be initiated. The monkeys received a liquid reward for releasing the bar 150 – 750 ms after target presentation. Target location was indicated by a cue (green square, second screen). Monkeys had to ignore any uncued event. (B) Recording sites. On each session, 24-contacts recording probes were placed in each FEF. (C) Behavioral performance of monkeys M1 and M2 at detecting the target in the presence or absence of a distractor (median % correct +/− median absolute deviations, dot correspond to individual sessions).

### Behavioral performance varies rhythmically at 4 to 7 cycles per hour

Hit and miss trials were sorted according to their respective time in the session and a running estimate of monkeys’ behavioral performance was estimated across the whole session time (see methods for detailed behavioral performance estimation procedures). Fig. 2a represents the running estimate of behavioral performance in an exemplar session, defined by the ratio between hit trials and the sum of hit and miss trials. In this figure, we observed a clear periodic fluctuation pattern of the behavioral performance by up to 30% along the entire session. The spectral density decomposition of this behavioral performance time series as estimated using wavelet transforms exhibits an oscillatory peak at 5.7 cycles per hour (Fig. 2g, gray, permutation test, p<0.05). Such rhythmic modulations of behavioral performance can be described consistently in all recording sessions and both subjects, at an average frequency of 5.59+ −0.30 cycles per hour (Fig. 2h, gray). Over all sessions, behavioral performance was, on average, 16% higher at the optimal phase (i.e. the phase maximizing behavioral performance) of each cycle of the behavioral oscillation compared to the anti-optimal behavioral phase (i.e. the phase minimizing behavioral performance, Fig. 2b, see methods for more details about average optimal phase to performance computations).

**Figure 2:**
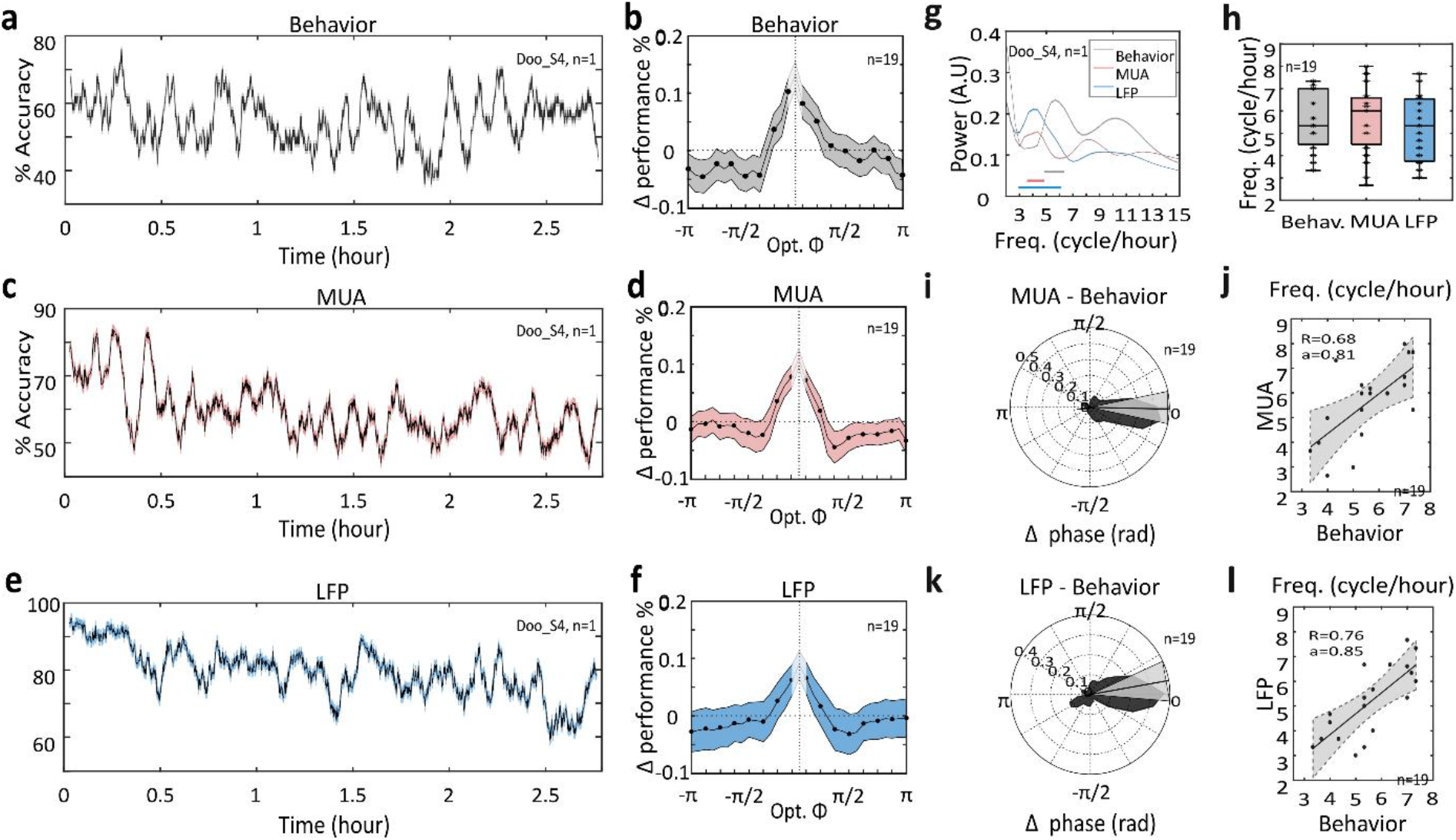
Ultra-slow rhythmic variations in behavioral performance and accuracies at decoding hit vs. miss trials from prefrontal MUA and LFP signals. (A) Running behavioral performance (Hits/(Hits+Misses)) estimates throughout session time (mean +/− s.e.) for the exemplar session *Doo_S4.* (B) Variations in overall behavioral performance across all sessions, as a function of phase difference from optimal phase. (C) Running estimates of target encoding classification accuracies for hit vs. miss classification from prefrontal MUA frequency content, throughout session time (mean +/− s.e.), for the same exemplar session as in (A). (D) Variations in overall hit vs. miss target decoding accuracies from prefrontal MUA across all sessions, as a function of phase difference from optimal phase. (E) Running estimates of target encoding classification accuracies for hit vs. miss classification from prefrontal LFP frequency content, throughout session time (mean +/− s.e.), for the same exemplar session as in (A). (F) Variations in overall hit vs. miss target decoding accuracies from prefrontal LFP frequency content across all sessions, as a function of phase difference from optimal phase. (G) Frequency power spectrum of the time series presented in (A, grey), (C, pink) and (E, blue). Epochs of statistical significance against the 95% C.I. indicated by lines of same color. (H) Distribution of ultra-slow oscillatory peaks across all sessions and both subjects (median +/− s.e.), for behavioral performance (grey) and hit vs. miss decoding accuracies from prefrontal MUA (pink) and LFP (blue), all identified peak frequencies were significant against 95% C.I (see method). (I-K) Phase difference and (J-L) correlation between ultra-slow oscillations identified in behavioral performance and MUA or LFP hit vs. miss decoding accuracies.

Previous studies indicate that variations in behavioral performance might be interpreted as fluctuations in how accurately subjects encode the task-relevant visual information(Busch et al., 2009; Jasper et al., 2019; Parto Dezfouli et al., 2018). Accordingly, an enhanced (respectively impaired) prefrontal representation of visual stimuli is expected during the phase of optimal (respectively poor) behavioral performances (van Vugt et al., 2018). In order to test this hypothesis, we apply machine learning procedures(Astrand et al., 2016, 2020; Gaillard et al., 2020a) to ongoing LFP/MUA activities following target presentation to decode behavioral outcome (Hit or Miss) based on FEF neuronal response to target presentation. Specifically, we analyze how much the ultra-slow oscillations detected in behavioral performances impact correct (Hit) or incorrect (Miss) response prediction based on FEF neuronal population responses to the target (See methods for more details about decoding procedures).

Figure 2c represents, for the same session as presented in figure 2a, the running estimates of decoding accuracies for hit vs. miss classification from prefrontal MUA frequency content, throughout session time. We observe that decoding accuracy varies in time by up to 34% on this exemplar session (Fig. 2c). A clear rhythmic fluctuation of decoding accuracies can be identified, peaking at 4.74 cycles per hour (Fig. 2g, pink, permutation test, p<0.05), closely matching the frequency identified in the behavioral performance (5.7 cycles per hour). Rhythmic modulation of decoding accuracies can be described consistently in all recording sessions from both subjects at an average frequency of 5.67 +−0.27 cycles per hour (Fig. 2h, pink). Over all recording sessions, decoding accuracies are on average over 13% higher at the optimal behavioral phase (i.e. the phase maximizing decoding accuracy) as compared to the anti-optimal behavioral phase (i.e. the phase minimizing decoding accuracy, Fig. 2d). Similarly, we observe rhythmic modulations of LFP based behavioral classification (Fig. 2e, blue) identified for this exemplar session at a specific rhythm of 4.6 cycles per hour. This rhythm was identified in all sessions at an average peak frequency of 5.18+-0.33 cycles per hour (Fig. 2h, blue). Over all recording sessions, decoding accuracies were, on average, over 11% higher at the optimal behavioral phase as compared to the anti-optimal behavioral phase (Fig. 2f).

Crucially, across all recording sessions, peak oscillatory frequencies identified in the behavioral performance and in the MUA decoding accuracy of hits vs. misses are highly correlated (Fig. 2j, Pearson correlation, r^2^=0.68, p <0.001). This is remarkable given the fact that at these time scales, any source of noise is expected to have a strong impact on the identification of peak frequency from any of the three considered time series. In addition, a clear phase locking between these two rhythms can be observed between the two signals (Fig. 2i, circular Kuiper’s test for equal distributions, p <0.05), confirming that the identified periods of low (resp. high) subject behavioral performance coincide with decreased (resp. increased) prefrontal accuracy of visual stimuli coding. Similarly, the frequency peaks of the LFP based target decoding accuracies highly correlate with the behavioral oscillatory peaks (Fig. 2i, Pearson correlation, r^2^=0.76, p<0.001), and are highly significantly phase locked (Fig. 2k, circular Kuiper’s test for equal distributions, p<0.05). Complementing these data, the peak frequency of the LFP based target decoding accuracies highly correlate with the MUA based target decoding accuracies oscillatory peaks (Supplementary Fig. 2, Pearson correlation, r^2^=0.63, p <0.01), and are significantly phase locked (Supplementary Fig. 2, circular Kuiper’s test for equal distributions, p <0.05).

Overall, these results indicate a functional link between the observed change in discriminability between hits and misses from the MUA/LFP signals and behavior. Behavioral performance on this specific attentional task is dependent of the correct allocation of attentional processes (Bello et al., 2020; Posner, 2016). As a result, the ultra-slow fluctuations in target processing and perception is expected to directly impact FEF attentional-related information(Astrand et al., 2016; Gaillard et al., 2020a) to the same extent.

### Prefrontal MUA based decoded attentional information varies rhythmically at 4 to 7 cycles per hour

Prefrontal population attention related information, can be assessed by machine learning techniques applied to ongoing prefrontal population multiunit (MUA) neuronal ensemble activity(Astrand et al., 2016; De Sousa et al., 2021; Gaillard et al., 2020a). In the following, we show that prefrontal population attention-related information is modulated at an ultra-slow rhythm that correlates with the rhythmic variations in behavioral performance described above. Specifically, we focus on correct trials, i.e. on trials in which spatial attention was (a priori) correctly oriented to the expected target location. On each correct trial, we compute the locus of the attentional spotlight just prior to target presentation and we test whether attention was properly oriented in the cued visual quadrant or not. We then compute a running average of attention decoding accuracy throughout the time of the session (Fig. 3a, same exemplar session as in figure 2a and 2c). We observe that decoding accuracy varies in time by more than 20%. In other words, how well attention-related information can be extracted from the prefrontal cortex population activity varies as a function of time in the session. Using wavelet transform, we identify for this specific session a peak in MUA attention-related information at 5.67 cycles per hour, matching the peak identified in the behavioral performance time series (Fig. 3e). Such rhythmic modulations in prefrontal population attention information can be described consistently in all recording sessions and both subjects, at an average frequency of 5.55+−0.23 cycles per hour (Fig. 3f, magenta). On average, over all recording sessions, MUA-based attention related information estimates were over 10% higher at the optimal phase as compared to the anti-optimal phase (Fig. 3b). Crucially, ultra-slow rhythmic variations in prefrontal attention-related information are highly correlated with the rhythmic variations in behavioral performance in the same sessions (supplementary Fig. 3a, Pearson correlation, r^2^=0.72, p<0.01).

**Figure 3:**
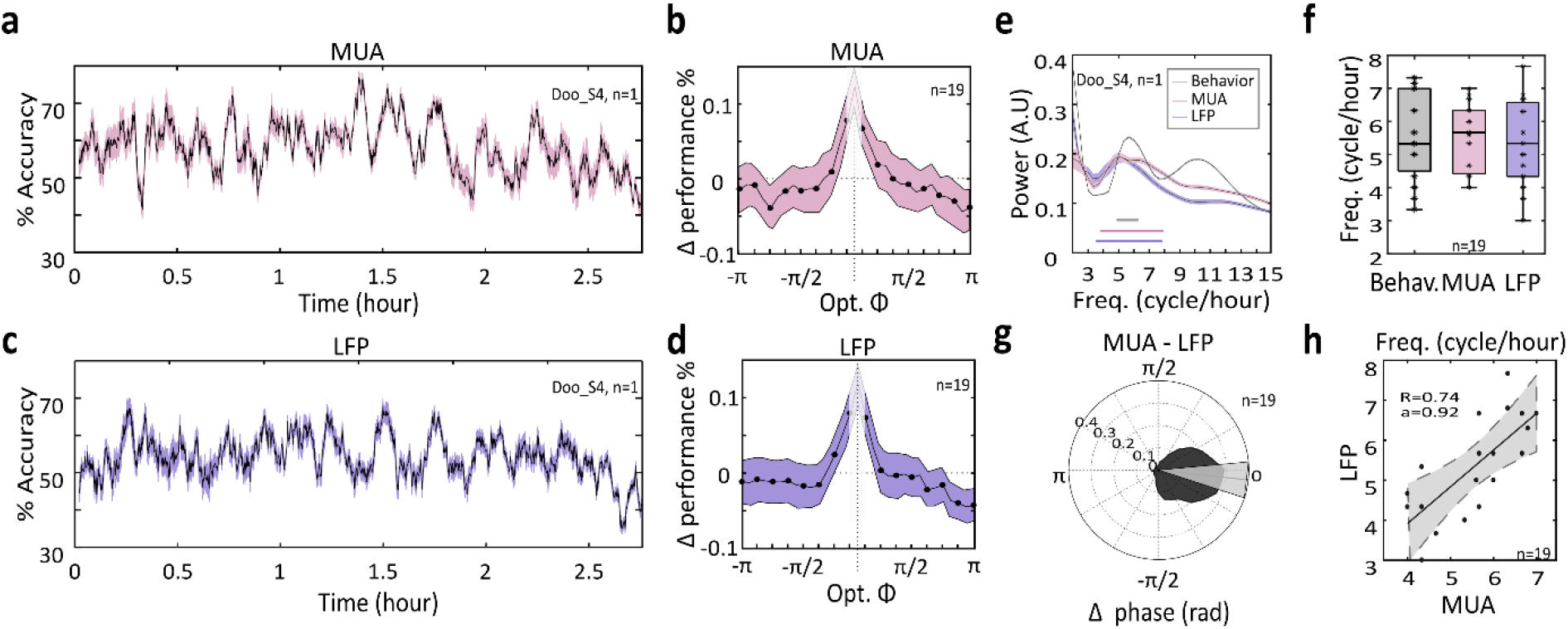
Ultra-slow rhythmic variations in prefrontal MUA and LFP attention-related information. (A) Running estimates of attention decoding accuracies from prefrontal MUA frequency content, throughout session time (mean +/− s.e.), for the exemplar session *Doo_S4* (same as in figure 2). (B) Variations in overall MUA-based attention decoding accuracies across all sessions, as a function of phase difference from optimal phase. (C) Running estimates of attention decoding accuracies from prefrontal LFP frequency content, throughout session time (mean +/− s.e.), for the same exemplar session as in (A). (D) Variations in overall LFP-based attention decoding accuracies across all sessions, as a function of phase difference from optimal phase. (E) Frequency power spectrum of the time series presented in (A), (magenta) and (B), (purple). (F) Distribution of ultra-slow oscillatory peaks across all sessions and both subjects (median +/− s.e.), for MUA-based (magenta) and LFP-based (purple) attention decoding accuracies. (G) Phase difference and (H) correlation between ultra-slow oscillations identified in the MUA-based and LFP-based attention decoding accuracy time series.

Critically, subjects show more precise attentional orientation during the good phase of the ultra-slow oscillation. Indeed, the rhythmic fluctuations in MUA attention-related information correspond to variations in the distribution of the attentional spotlight prior to target presentation on the trials falling on the peak (Fig. 4a) relative to those falling on the troughs of the ultra-slow rhythmic variations (Fig. 4b). Specifically, attention is more focused and closer to the expected target location by an average of 2.1°, on peak trials relative to trough trials (difference in peaks of hit maps). The net effect of this is a covert attentional exploration of the cued location during the peak trials and a more distributed exploration of the screen on trough trials (Fig. 4c). These changes in prefrontal MUA attention-related information do not coincide with systematic changes in MUA average attention-related responses prior to target presentation. Indeed, there is no significant systematic variation in average MUA responses on peak trials relative to trough trials (Fig. S4 Wilcoxon rank sum test, p>0.5). This confirms that the change in overall MUA attention-related information does not correspond to a non-specific mechanism such as metabolic depletion, but rather to a rhythmic change in the prefrontal neuronal representation of attentional processes.

**Figure 4:**
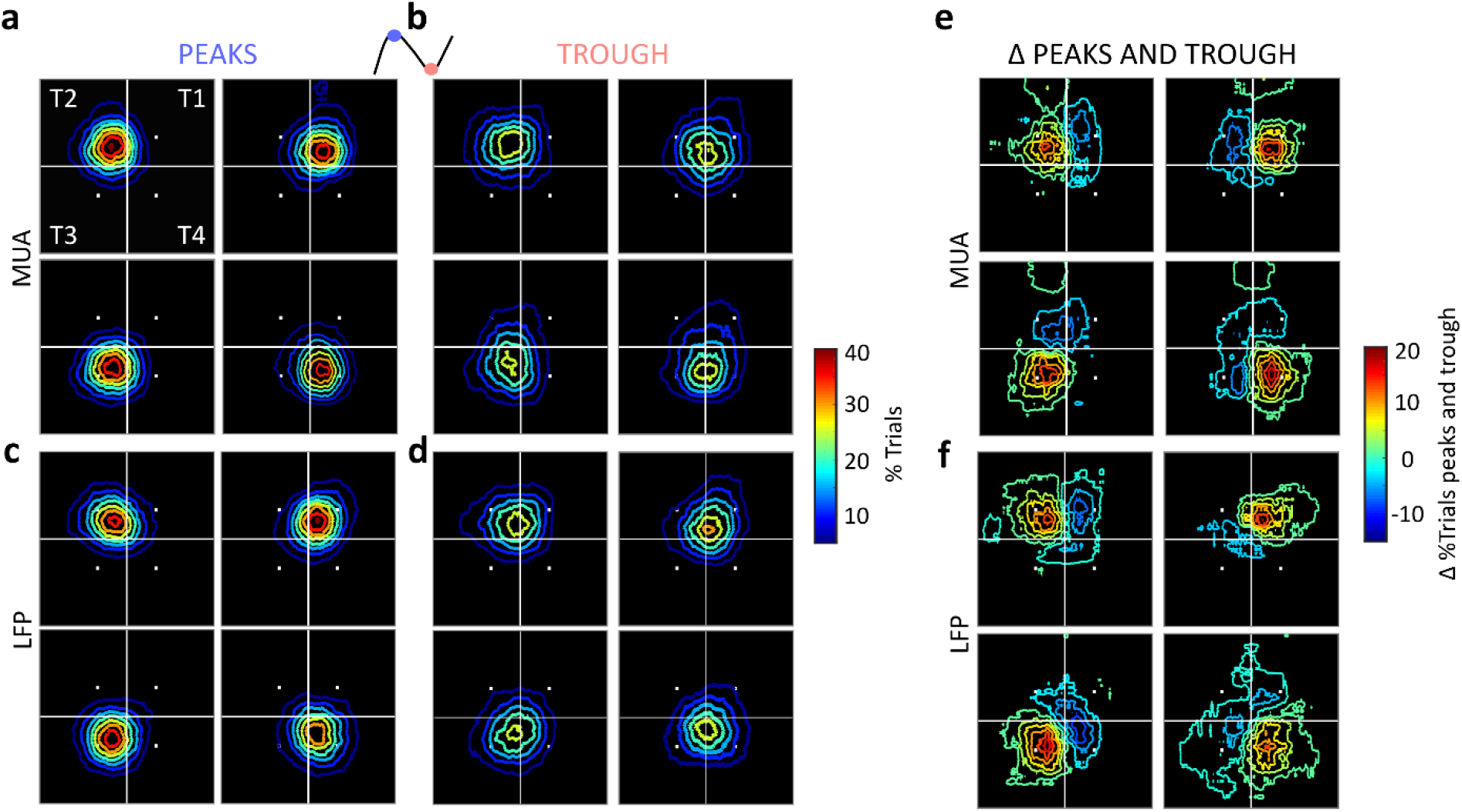
Attention-related information in the MUA (upperpanel) andLFP (lowerpanel)signals is more focused and closer the cued location on peaks of ultra-slow oscillations (A, C, resp.) as compared to troughs (B, D, resp.). In each panel A to F, each black square represents the screen, the vertices of the central gray square indicate the locations where the cues are presented. The colored traces in panels A to B represent density (% of trials) plots of where the spotlight falls on the screen, prior to target presentation, on correct trials, when the attention is cued to the upper left quadrant (upper left square), upper right (upper right square), lower right (lower right square) and lower left square (lower left square, see cartoon on the upper right). Panels E and F represent the difference between the maps presented in A and B, and C and D respectively.

### Prefrontal LFP based decoded attentional information varies rhythmically at 4 to 7 cycles per hour

These variations in the population MUA attention-related information reflect changes in the population neuronal code that subserves spatial attention in the prefrontal cortex. In order to test whether these ultra-slow rhythmic fluctuations in the attentional code correspond to a local mechanism or whether they reflect on more mesoscopic signals such as local field potentials (LFP), we reproduced the previous analysis on these latter signals. Indeed, previous studies have demonstrated that very much like for MUA signals, attention can be classified from the single trial LFP power content (mostly from the gamma band), though to a lesser extent than from MUA signals(De Sousa et al., 2021; Tremblay et al., 2015). On each correct trial, we compute the locus of the attentional spotlight just prior to target presentation based on the recorded prefrontal LFP signals and we test whether attention was properly oriented in the cued visual quadrant or not. We then compute a running average of attention decoding accuracy throughout the time of the session (Fig. 3c, same exemplar session as in figure 2a, 2c, 2e and 3a). As was seen for behavioral performance, hit/miss decoding from prefrontal LFPs and MUA-based attention decoding, we observe that decoding accuracy varies in time by up to 20%. Using wavelet transform, we identify for this specific session a peak in LFP spatial related information at 5Hz, matching the peak identified in the behavioral performance and MUA based attention information time series (Fig. 3e). Such rhythmic modulations in prefrontal LFP attention information can be described consistently in all recording sessions and both subjects, at an average frequency of 5.34 +-0.29 cycles per hour (Fig. 3f, purple). On average, over all recording sessions, LFP-based attention related information estimates are over 10% higher at the optimal phase as compared to away from this optimal phase (Fig. 3d). Crucially, as is observed for MUA-based attention information, ultra-slow rhythmic variations in prefrontal attention related information are correlated with the rhythmic variations in behavioral performance in the same sessions (supplementary Fig. 3b, Pearson correlation, r^2^=0.58, p<0.05). Interestingly, a significant correlation is observed across sessions between the oscillatory peaks of the MUA based attention related information and the LFP based attention related information. (Fig. 3g, r^2^=0.73, p<0.001). In addition, we report a significant phase locking between the MUA and LFP ultra-slow oscillations in attention-related information across all sessions (Fig. 3g, circular Kuiper’s test for equal distributions, p <=0.05), suggesting a common origin or mechanism.

As observed for the MUA-based attention-related information, the rhythmic variations in LFP attention-related information coincide with an over attentional exploration of the cued location during the peak trials (Fig. 4c) and a more distributed exploration of the screen on trough trials (Fig. 4d and 4f, 3.6° average difference in peaks of hit map between the two conditions).

### MUA and LFP markers of the attentional ultra-slow oscillations

Attention-related information modulation by ultra-slow oscillations could be observed from both LFP and MUA signals. This suggests that the observed slow oscillations might not reflect a local process but rather a more global mesoscopic process mediated by LFPs. In order to test this hypothesis, we computed prefrontal LFP power spectra during the 1000ms attentional period preceding target presentation, on correct trials, using a continuous wavelet transform (see methods), and we segregated individual trial measures into trials falling in the peak or trough of the ultra-slow oscillations of the MUA attention-related information (see methods for more details). Average power spectrum differences across all sessions (n=19), all recording channels (n=48 per session) and the two monkeys reveal a significant higher power in the peak relative to the trough of the oscillations, in two specific functional frequency bands (Fig. 5a, significance assessed against 95% C.I.): a theta frequency band (mean= 6.35 Hz +−1.65 s.e.), and a beta frequency band (mean= 24.97 Hz +−3.58 s.e.). Importantly, theta and beta modulation of LFP activities have been associated with orientation of attention (Buschman and Miller, 2007b; Fiebelkorn et al., 2018a; Gaillard et al., 2020a; VanRullen, 2018). The observed modulation of these two frequency ranges during the ultra-slow oscillation cycles supports the idea that these oscillations specifically impact spatial attention neuronal processes, and thereby behavioral performance.

**Figure 5:**
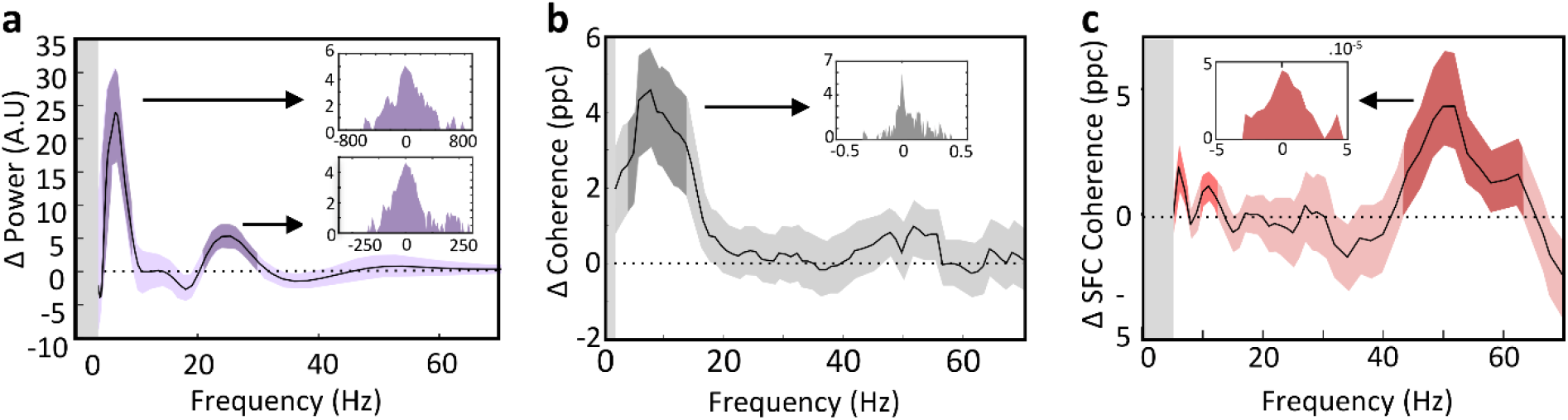
Ultra slow oscillations in MUA and LFP attention-related content coincides with significant changes in LFP power (A), LFP coherency (B) and spike field coherency (C). All metrics are extracted over the 1000ms preceding target presentation. For each metric, the main plot represents the difference between the metric obtained for trials at the peak of the slow oscillations vs. trials at the trough of the slow oscillations, averaged over all recording signals (n=912 channels, mean +− s.e., above 95 C.I in darker shades). Insets represent the distribution of the modulation index of each metric between the peak and trough of the slow oscillation trials, averaged over significance epoch (black arrow), over all recorded signals (n=912 channels).

In addition, we observe during the same trial epoch (1000ms before target presentation), enhanced coherency across LFPs, in the correct trials located at the peak relative to the trough of the ultra-slow oscillations, as assessed from a coherency measure (pairwise phase consistency). This enhanced coherency is specifically observed in the alpha range (average peak=9.25 Hz +−7.35 s.e., Fig. 5b, significance assessed against 95% C.I.). The majority of LFP pairs are up-modulated in the peak relative to the trough of the oscillation cycles (Figure 5b, inset). This suggests that the FEF receives long-range signals, modulating overall alpha coherency of the LFPs all across the FEF map, bilaterally.

We further show that spike field-coherency estimated during the same trial epochs as LFP power and coherency, are enhanced on the correct trials located at the peak relative to the trough of the ultraslow oscillations in three specific frequency ranges (Fig. 5c, significance assessed against 95% C.I.): the theta (average peak=6.25 Hz+−0.75 s.e.), alpha (average peak=11 Hz +−1.30 s.e.), and gamma range (average peak=50.75 Hz +−6.75 s.e.). Changes in spike-field coherence in the gamma band are reliably associated with changes in attentional processes orientation(Chalk et al., 2010; Fiebelkorn et al., 2019; Fries, 2015a; Fries et al., 2001).

Overall, these results indicate that the observed ultra-slow fluctuations in MUA-related attentional processes are subsequent to long-range changes impacting prefrontal LFPs, and are locally mediated through specific changes in spike-field coherence mechanisms.

### Human behavioral performance and pupil size fluctuates at 4 to 7 cycles per hour

In order to test whether the ultra-slow rhythm described above is specific to high attentional demand tasks, we had 30 human subjects detect a shivering Gabor in the midst of slowly drifting Gabors of variable orientations for 75 minutes with no interruption. Figure 6a represents the running estimate of an exemplar subject detection times (i.e. time taken by the subject to identify a new shivering Gabor) across a single session. As seen in the behavioral performance of monkeys, the detection times fluctuate by up to 15 seconds. A wavelet analysis reveals a significant oscillatory peak at ~ 5.85 cycles per hour (Fig. 6b, permutation test, p<0.05). This specific rhythm in subjects’ detection time was consistently identified across subjects, at an average frequency of 5.96 +−0.25 cycles per hour (Fig. 6e, orange, mean +− s.e., n=35).

**Figure 6:**
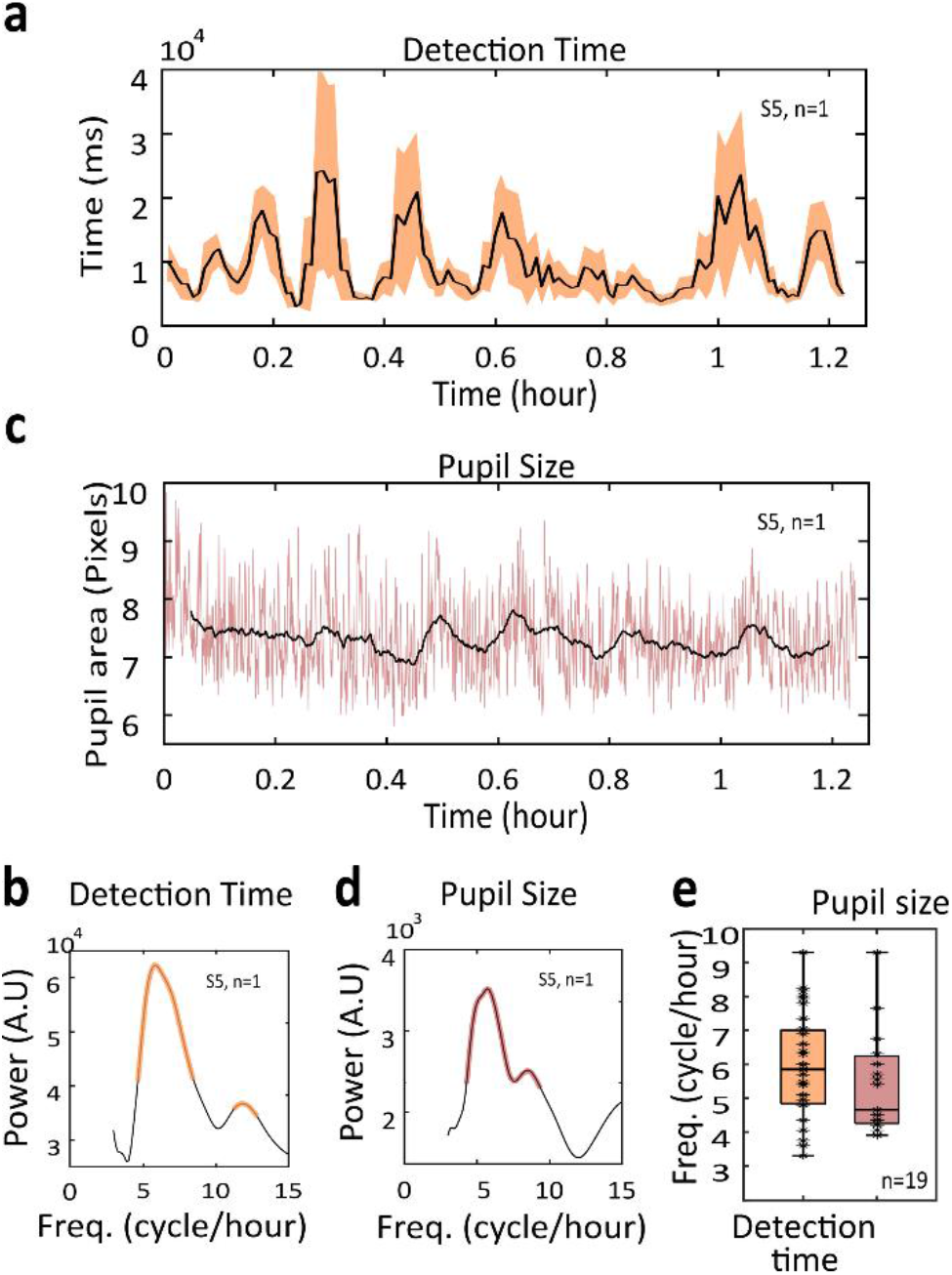
Ultra-slow rhythmic fluctuations in behavioral performance and pupil size in human subjects during sustained cognitive activity. (a) Detection time estimates throughout session time (mean +/-s.e.) for the exemplar subject S5 and (b) corresponding power spectrum (orange, test for random permutations, p<0.05). (c) Pupil area estimates throughout session time (black line: convoluted pupil size, pink line: raw pupil size data) for the exemplar subject S5 and (d) corresponding power spectrum (pink, test for random permutations, p<0.05). (e) Distribution of ultra-slow oscillatory peaks across all subjects (median +/− s.e.), for detection times (orange) and corresponding pupil size (pink), all frequency peaks significant in all subjects (random permutation p<0.05).

Changes in pupil size is associated with a broad range of cognitive processes(Beatty and Lucero-Wagoner, 2000; Sirois and Brisson, 2014; van der Wel and van Steenbergen, 2018), including attention(B et al., 2012), and working memory(Just et al., 2003; Kramer, 1990). In addition, increase in pupil size has been taken as an indirect index of the noradrenergic function(Dragone et al., 2018; Reynaud et al., 2019). Accordingly, in a subset of subjects (n=19), we recorded pupil size while performing the task. Figure 6b represents the running estimate of an exemplar subject pupil size across a recording session (same exemplar session as Fig. 6a). A wavelet analysis reveals a significant frequency peak at 5.7 cycles per hour (Fig. 6d, permutation test, p<0.05), consistently identified across subjects, at an average frequency of 5.56 +− 0.43 cycles per hour (Fig. 6e, pink, mean +− s.e., n=19). Detection times and pupil sizes peak frequency distributions across subjects were highly similar (Ranksum test for equal mean, p>0.57, n=19).

## Discussion

Overall, we show that when performing an attentional task for one hour or more, the behavior of both humans and non-human primates consistently fluctuates between periods of high and low performance at a rhythm of 4 to 7 cycles per hour (every 9 to 15 minutes). These behavioral fluctuations are observed in both a difficult (in non-human primates) and an easy (in humans) attentional task, indicating that these fluctuations are not dependent on the task demand. At the physiological level, these epochs of high and low performance coincide with fluctuations in pupil diameter, indicating underlying changes in arousal and information-processing load. Accordingly, we show that these behavioral fluctuations coincide, at the neurophysiological level, with fluctuations in the attention and perception coding accuracy of prefrontal neuronal populations. We further identify specific markers of these fluctuations in LFP power, LFP coherence and spike-field coherence, suggesting long-range rhythmic modulatory inputs to the prefrontal cortex rather than a local prefrontal origin.

### Novel ultra-slow rhythmic fluctuations in behavior and neuronal processes

Ongoing fluctuations in neural activity that are independent of sensory processes have been described in the brain at multiple time scales(Arieli et al., 1996; Aston-Jones and Cohen, 2005; Beaman et al., 2017; Cowley et al., 2020; Engel et al., 2016; Gaillard et al., 2020b; McGinley et al., 2015, 2015, 2015; Milton et al., 2020; Oby et al., 2019; Okun et al., 2015, 2019; Reimer et al., 2014; Stringer et al., 2019; Vinck et al., 2015; Wasmuht et al., 2019). These fluctuations are shown to correlate with behavioral and physiological variables such as pupil diameter(Ebitz and Moore, 2017; Gutnisky et al., 2017; Joshi et al., 2016; McGinley et al., 2015; Nassar et al., 2012; Okun et al., 2015; Stringer et al., 2019; Vinck et al., 2015), eye movements(Leopold and Logothetis, 1998; Steinmetz and Moore, 2014), wakefulness(McGinley et al., 2015; Milton et al., 2020; Reimer et al., 2014), or attention(Engel et al., 2016; Huang and Elhilali; Moore and Zirnsak, 2017; Rabinowitz et al., 2015; Snyder et al., 2016).

Some of these fluctuations in behavior and neuronal processes are rhythmic, spanning a wide range of frequencies from 0.01Hz(Chan et al., 2017; Klein and Armitage, 1979; Palva and Palva, 2011), up to 100Hz (100 cycles per second(Fries, 2005; Jia and Kohn, 2011)). Attentional and perceptual cortical processes have specifically been associated with theta frequencies (3-6 Hz(Dugué et al., 2014, 2016; Fiebelkorn et al., 2018a) as well as alpha (7-12 Hz(Gaillard et al., 2020b)), beta (18-25 Hz(Hassen et al., 2019; Vernet et al., 2019)) and gamma frequencies (25-100 Hz(Fries, 2015a)). For example, in the primate primary visual cortex, Engel et al. (2016) describe an alternation of on and off spiking episodes of 149ms and 100ms average duration respectively(Engel et al., 2016). This pattern of alternation is similar to that described during sleep or anesthesia. More recently, Gaillard et al., (2020) describe rhythmic fluctuations in prefrontal attention-related neuronal activity in the 7-12 Hz range(Gaillard et al., 2020b). McGinley et al (2015) show that cortical membrane potential in the mice auditory cortex co-varies with pupil size at two different signature frequencies of 0.03 Hz and 0.3 Hz respectively, as a function of general arousal(McGinley et al., 2015). All of these variations are predictive of optimal neuronal and behavioral processes, but all happen at a much faster time scale than the one described in the present study. A recent study describes widespread changes in neuronal dynamics in the dlPFC of the freely moving macaque monkey(Milton et al., 2020). This study identifies transitions between desynchronized neuronal states during active wakefulness and synchronized neuronal states during quiet wakefulness. In contrast with our observations, synchronous activity was associated with a significant decrease in spiking rates and an increase in the power of the slow LFP oscillations in the 0.5-10Hz range. These transitions were highly correlated with spontaneous motion and took place every 30 minutes to 60 minutes, thus at a slower rate than described here. At closer inspection of the Milton et al. data(Milton et al., 2020), however, faster transitions can be seen, predominantly in the wake state, at the same rhythm as described here. This suggests that the rhythmic fluctuations we describe interact with wake/rest transitions but stem from an independent neuronal system mechanism. This will have to be further explored experimentally.

Up to now, the description of slow rhythms in the range of the hour has been missing. This is due to the fact that data is often collected in short blocks of 5 to 20 minutes and subjects are allowed to rest in between, thus disrupting the temporal structure in sustained workload processes at each pause. By recording behavioral performance during sessions lasting from 1 to 4 hours (with no externally generated interruptions), we report rhythmic fluctuations in behavioral performance in the range of 4 to 7 cycles per hour. These fluctuations are consistent across individuals and species. In addition, they are independent from task attentional demand (as they can be reported in both a low and a high attentional demand task) and subjects’ expertise (as they can be reported both in naïve human subjects and over-trained macaques). Importantly, they occur in the absence of any kind of monotonic change in behavioral performance as time goes by, in contrast with what has been described by others(Cowley et al., 2020). This thus goes against current theories hypothesizing a decrement of performance under sustained-attention constraints (e.g. the resource-control hypothesis(Thomson et al., 2015)). Over longer periods of days and even weeks, mental effort and cognitive fatigue have been shown to progressively degrade cognitive performance(Lockley et al., 2004; Proctor et al., 1996; Virtanen et al., 2009). It will be crucial to understand how this decay in cognitive performance interacts with the behavioral ultra-slow rhythms we describe here and whether there is a functional link between these two observations.

### Neuronal correlates of the ultra-slow fluctuations in behavioral performance

We specifically show that both prefrontal attentional and perceptual neuronal processes are altered during the ultra-slow rhythmic fluctuations, directly impacting behavioral performance. The net effect of the described rhythmic fluctuations is a higher attentional and perceptual information content, i.e. neuronal processes closer to optimality, during high behavioral performance epochs compared to low performance epochs. Importantly, these changes take place in the absence of significant overall changes in neuronal spiking rate. They thus are not associated with a global variation in neuronal population excitability(Milton et al., 2020), but rather with a change in overall population informational coding accuracy.

We find that these rhythmic modulations of behavior also impact several neuronal markers that have been associated with attentional and perceptual processes. First, we report an enhancement of the power of prefrontal LFP activity in the alpha(Bollimunta et al., 2008; Gaillard et al., 2020a; Mo et al., 2011) (6-9Hz) and beta(Buschman and Miller, 2009; Fiebelkorn et al., 2018b; Siegel et al., 2008) frequencies (20-30Hz). Second, we show that on high behavioral performance epochs, LFP alpha coherency is higher than on low behavioral performance epochs. Signal coherency within and between brain areas is proposed as a key mechanism at the core of brain computing functions and supports information/stimulus processing(Fries, 2015b), accounting for network connectivity(Vinck et al., 2010), and changes in behavioral performance(Zareian et al., 2018, 2020). Last, we report changes in spike field coherency within the prefrontal cortex between the epochs of high vs. low behavioral performance, in the same frequency ranges as reported by others during attention orientation(Fries et al., 2008; Gregoriou et al., 2009b; Paneri and Gregoriou, 2017), thus pointing to a direct modulation of attentional mechanisms.

Overall, this supports the idea that epochs of high vs low behavioral performance coincide with specific changes in the attention and perception processes. These changes arise from long-range mechanisms supported by LFP coherency and LFP power content in specific frequency bands repeatedly associated with attentional processes, as well as from local changes in spike-field coherence in these very same frequency bands. The origin of these fluctuations, whether arising from global brain network configuration, or from a specific brain region controller, remains to be explored. It is unclear whether the observed very slow rhythmic processes only impact the functions being recruited by the task, namely attention and perception (and hence specifically the functional brain network subserving this function), or whether theses fluctuations impact all cognitive performances, irrespective of the task. Again, this will have to be further explored.

### Relationship with attentional lapses and mind wandering

During the execution of cognitive tasks in humans, “task-off” or “mind-wondering” states (in contraposition to “task-on” periods) can be observed(Compton et al., 2019; Smallwood and Schooler, 2015). These “task-off” epochs have been associated with specific neural signatures such as increased scalp alpha activity(van Dijk et al., 2008) and an enhanced activation of the default-mode-network (Knyazev et al., 2011; Mo et al., 2013). These states of attentional lapses have been explained in the context of resource-control theory(Thomson et al., 2015), that posits that (i) an executive control is required to sustain active goal maintenance, (ii) the amount of attentional resources available for any given individual is fixed and (iii) attentional resources are consumed both during task-on and task-off states. As a consequence, this model predicts that executive control wanes over task-on periods, resulting in performance-costs in task-off periods.

Our data show oscillations in behavioral performance. These behavioral oscillations coincide with fluctuations in prefrontal attentional coding accuracy (or resources). As a result, they can be seen as alternations between task-on and task-off periods. However, attentional resources alternate between states of depletion (transitory waning) and states of efficient processing (restoration), instead of waning with time, as initially predicted by the resource control theory. These fluctuations in behavior also coincide with fluctuations in pupil size. This physiological parameter has been shown to depend on the noradrenergic system(Dragone et al., 2018; Reynaud et al., 2019) and to be altered in patients with attention disabilities(Beane and Marrocco, 2004). The rhythmic transitions between task-on and task-off epochs and associated attentional and perceptual processes could thus be under the direct influence of the Locus Coeruleus and noradrenergic neuromodulation. This will have to be further explored.

### Possible origin of ultra-slow rhythmic fluctuations

Changes in brain states have been reported in the wake state and during the different sleep cycles or during states of altered consciousness such as anesthesia(Barttfeld et al., 2015; Demertzi et al., 2019). None of these changes take place at frequencies nor coincide with as specific neuronal correlates in LFP power, LFP coherency and spike field coupling as described in the present work. Thus, the mechanistic origin of the behavioral and neurophysiological rhythmic fluctuations reported in this manuscript remains elusive. One possible explanation is that these fluctuations arise as a consequence of cognitive fatigue, in which one can expect a partial impairment of the neuronal processes supporting the attentional function, thus limiting behavioral performance by decreasing alertness and information processing. This hypothesis would predict alternations of metabolic depletion and regeneration, and a specific impact on the recruited functional network rather than on the entire brain. A second explanation for this alternation between optimal and suboptimal cognitive abilities could be an evolutionary optimization strategy. Runners on very long trails must adapt their speed, alternating between periods of sprints and slower running periods, so as to maintain an overall performance and accomplish an optimal global timing. If they run too fast too often, they risk to collapse out of exhaustion. If they run too slow, they will not maximize their reward, be it a prize or an individual record. In a similar way, ultra-slow oscillations can allow to maintain an acceptable level of cognitive performance across very long periods of time without depleting the entire brain cognitive resources, which would eventually result in a complete shutdown of all processes. A third hypothesis is that multilevel oscillatory activity could arise from the specific organization of the primate brain. Specifically, many cognitive functions often rely on complex neuronal networks involving multiple areas related to each other by complex feedforward and feedback connections. Indeed, reports show that synchronized brain rhythms could arise in densely wired systems (such as the primate brain) in a small-world way(Kim and Lim, 2013; Li et al., 2015) and propose that a scale-free network structure of the brain could be responsible for large scale rhythmic activity (Mi et al., 2013). Such multi-scaled and hubs organized structures are described to be highly adapted against local impairment or lesion. Modelling work would help support or infirm this possibility. A fourth hypothesis would be a neuromodulatory origin. The locus coeruleus, which distributes norepinephrine all across the brain through widespread projections (Aston-Jones and Cohen, 2005) modulates arousal. In particular, its neuronal activity has been linked to variability in pupil size(Joshi et al., 2016; Liu et al., 2017; McGinley et al., 2015) and behavioral performance(Aston-Jones and Cohen, 2005; Eldar et al., 2013). Another candidate is acetylcholine, which is released by the basal forebrain, and impacts pupil size as well as locomotion(Everitt and Robbins, 1997; McCormick, 1992; McGinley et al., 2015; Yüzgeç et al., 2018). These multiple hypotheses will have to be evaluated experimentally.

### Ultra-slow rhythms and attentional reallocation

Tradeoff between exploratory and exploitation is crucial for optimal behavior. For example, a bee can keep collecting pollen from the same flower (exploitation), which comes at a cost of lower harvest (all pollen will end up being collected), or move (exploration) from one flower to the next(Kembro et al., 2019) (exploration), which comes at a strong energetic cost (moving from one flower to the next). Allocation of human brain attentional mechanisms is proposed to alternate between exploration and exploitation strategies(Gaillard et al., 2020a; VanRullen, 2018). The rhythmic drop in the accuracy of ongoing cognitive processes that we report here might actually leave an open window for the subject to shift cognitive resources from the ongoing process to another process. From an ecological perspective, this would force on the subject a kind of cognitive flexibility, possibly originating from general large-scale transitions in brain functional networks. Such transitions have been described at lower time scales during normal brain states and states of altered consciousness(Huang et al., 2020).

### Conclusion

Modern societies and the rise of assistive technologies result in a whole new framework where human cognitive production is shifted from low cognitive demand tasks to sustained demanding cognitive tasks over long time periods. It is thus crucial to understand how the human brain is able to produce and sustain high demand cognitive processes efficiently, without producing and accumulating errors. In this context, the slow rhythmic behavioral and cognitive fluctuations described here are expected to instruct on how to optimize work and school environments, in a way that is more adapted to brain physiology. In addition, they are a promising targets for pharmacological or neuro-modulatory enhancement using behavioral training, transcranial magnetic stimulation or neurofeedback(Bagherzadeh et al., 2020; Dugué and VanRullen, 2017; Dugué et al., 2014; Horschig et al., 2015; Ros et al., 2017; Saj et al., 2018). These methods have up to now targeted infra-second rhythms. Applying them to the described ultraslow behavioral and cognitive rhythm is expected to open interesting perspectives in the field of human productivity, learning and teaching.

## Acknowledgments

S.B.H was supported by ERC Brain3.0 #681978, ANR-11-BSV4-0011 & ANR-14-ASTR-0011-01, LABEX CORTEX funding (ANR-11-LABX-0042) from the Université de Lyon, within the program Investissements d’Avenir (ANR-11-IDEX-0007) operated by the French National Research Agency (ANR). C.D.S., C.G., J.A. and C.L. were supported by ERC Brain3.0 #681978. We thank research engineer Serge Pinède for technical support and Jean-Luc Charieau and Fidji Francioly for animal care. All procedures were approved by the local animal care committee (C2EA42-13-02-0401-01) and the Ministry of research, in compliance with the European Community Council, Directive 2010/63/UE on Animal Care. All human experiments were performed in accordance with relevant guidelines and regulations (the declaration of Helsinski). All experimental protocols were approved by the CNRS (research institution acting as promotor) and the CPP-Sud-Est (acting as licensing committee). The project authorization is #ID RCB 2017-A03612-51.

## Author contributions

Conceptualization, C.G., C.D.S. and S.B.H.; Data Acquisition, S.B.H.H., F.D.B., J.A., C.L., C.Z.; Methodology, C.G., C.D.S and S.B.H.; Investigation, C.G., C.D.S. and S.B.H.; Writing – Original Draft, C.G., C.D.S and S.B.H.; Writing – Review & Editing, C.G., C.D.S., J.A. and S.B.H.; Funding Acquisition, S.B.H.; Supervision, S.B.H.

## Material and methods

### Monkey experiments – Ethical statement

All procedures were in compliance with the guidelines of European Community on animal care (Directive 2010/63/UE of the European Parliament and the Council of 22 September 2010 on the protection of animals used for scientific purposes) and authorized by the French Committee on the Ethics of Experiments in Animals (C2EA) CELYNE registered at the national level as C2EA number 42 (protocole C2EA42-13-02-0401-01).

### Monkey experiments – Subjects and surgical procedure

Two male rhesus monkeys (Macaca mulatta) weighing between 6-8 kg underwent a surgery during which they were implanted with two MRI compatible PEEK recording chambers placed over the left and the right FEF hemispheres respectively (figure 1A), as well as a head fixation post. Gas anesthesia was carried out using Vet-Flurane, 0.5 – 2% (Isofluranum 100%) following an induction with Zolétil 100 (Tiletamine at 50mg/ml, 15mg/kg and Zolazepam, at 50mg/ml, 15mg/kg). Post-surgery pain was controlled with a morphine pain-killer (Buprecare, buprenorphine at 0.3mg/ml, 0.01mg/kg), 3 injections at 6 hours interval (first injection at the beginning of the surgery) and a full antibiotic coverage was provided with Baytril 5% (a long action large spectrum antibiotic, Enrofloxacin 0.5mg/ml) at 2.5mg/kg, one injection during the surgery and thereafter one each day during 10 days. A 0.6mm isomorphic anatomical MRI scan was acquired post surgically on a 1.5T Siemens Sonata MRI scanner, while a high-contrast oil filled grid (mesh of holes at a resolution of 1mmx1mm) was placed in each recording chamber, in the same orientation as the final recording grid. This allowed a precise localization of the arcuate sulcus and surrounding gray matter underneath each of the recording chambers. The FEF was defined as the anterior bank of the arcuate sulcus and we specifically targeted those sites in which a significant visual and/or oculomotor activity was observed during a memory guided saccade task at 10 to 15° of eccentricity from the fixation point (figure 1A). In order to maximize task-related neuronal information at each of the 24-contacts of the recording probes, we only recorded from sites with task-related activity observed continuously over at least 3 mm of depth.

### Monkey experiments – Behavioral task and Experimental setup

The task is a 100% validity endogenous cued target detection task (fig 1a). The animals were placed in front of a PC monitor (1920×1200 pixels and a refresh rate of 60 HZ), at a distance of 57 cm, with their heads fixed. The stimuli presentation and behavioral responses were controlled using Presentation (Neurobehavioral systems^®^, https://www.neurobs.com/). To start a trial, the bar placed in front of the animal’s chair had to be held by the monkeys, thus interrupting an infrared beam. The onset of a central blue fixation cross (size 0.7°×0.7°) instructed the monkeys to maintain eye position inside a 2°×2° window, defined around the fixation cross. To avoid the abort of the ongoing trial, fixation had to be maintained throughout trial duration. Eye fixation was controlled thanks to a video eye tracker (Iscan™). Four gray square landmarks (LMs – size 0.5°×0.5°) were displayed, all throughout the trial, at the four corners of a 20°x20° hypothetical square centered onto the fixation cross. Thus, the four LMs (up-right, up-left, down-left, down-right) were placed at the same distance from the center of the screen having an eccentricity of 14° (absolute x- and y-deviation from the center of the screen of 10°). After a variable delay from fixation onset, ranging between 700 – 1200 ms, a small green square (cue – size 0.2°×0.2°) was presented, for 350 ms, close to the fixation cross (at 0.3°) in the direction of one of the LMs. Monkeys were rewarded for detecting a subtle change in luminosity of this cued LM. The change in target luminosity occurred unpredictably between 350 – 3300 ms from the cue off time. In order to receive a reward (drop of juice), the monkeys were required to release the bar in a limited time window (150 – 750 ms) after the target onset (Hit trial). In order to make sure that the monkeys did use the cue instruction, on half of the trials, distractors were presented during the cue to target interval. Two types of distractors could be presented: (i) uncued landmark distractor trials (33% of trials with distractor); these corresponded to a change in luminosity, identical to the awaited target luminosity change, and could take place equiprobably at any of the uncued LMs; (ii) workspace distractor trials (67% of trials with distractor); these corresponded to a small square presented randomly in the workspace defined by the four landmarks. The contrast of the square with respect to the background was the same as the contrast of the target against the LM; when presented at the same radial eccentricity as the LMs, the workspace distractor had the same size as the landmarks; for smaller eccentricities, the size of the workspace distractor was adjusted for cortical magnification such that it activated an equivalent cortical surface at all eccentricities. The monkeys had to ignore all of these distractors. Responding to any of them interrupted the trial. If the response occurred in the same response window as for correct detection trials (150 – 750 ms), the trial was counted as a false alarm (FA) trial. Failing to respond to the target (Miss) similarly aborted the ongoing trial. Overall, data was collected for 19 sessions (M1 10 Sessions, M2 9 Sessions). The behavioral performance of each animal on correct trials is presented in figure 1b, depending on whether distractors were present or not.

### Monkey experiments – Electrophysiological recording

Bilateral simultaneous recordings in the two frontal eye fields (FEF) were carried out using two 24 contacts Plexon U-probes (fig. 1b). The contacts had an interspacing distance of 250 μm. Neural data was acquired with the Plexon Omniplex^®^ neuronal data acquisition system. The data was amplified 400 times and digitized at 40,000 Hz. A threshold defining the multi-unit activity (MUA) was applied independently for each recording contact and each session before the actual task-related recordings started.

### Monkey experiments – Behavioral performance as a function of session time

Hit and Miss trials were compiled relative to their absolute time position in the session. Behavioral performance, defined as hits/(hits+misses) was computed on successive sets of 50 trials, corresponding to an average time period of 5.83 ± 1.1 min. This procedure was iterated over the entire length of the session in 1 trial steps. Time sampling variability (due to variability in trial length and reaction times) was compensated by assigning, to each running behavioral performance estimate, its corresponding Gaussian mean trial time relative to the session (σ=length of testing set/6, here σ=4.166). This procedure resulted in a time series representing running behavioral performance as a function of time in the session.

### Monkey experiments – Signal frequency analyses of behavioral performance time series

Behavioral performance time series were interpolated (cubic spline function) for each independent session. Across sessions, this resulted in a sampling frequency of 1538.4±232 samples/hour for an average session duration of 3.526 hours ± 0.6796. Specifically, median task duration was 3.94 hours for monkey M1 (s.e. = 0.66) and 2.99 hours for monkey M2 (s.e. = 0.20). These parameters allow the description of frequencies ranging from 0.29 ± 0.515 to 769 ± 116 cycles per hour. The spectral analysis of this time series was performed on detrended data using a Morlet Wavelet transform. Standard error represented on the power spectrum is calculated along the time dimension of the wavelet analysis. We established statistical significance by reshuffling trials presentation time across the session (1000 times) and generating a random distribution of signal spectral content. Significance was then computed by comparing each session power spectrum to its corresponding 95% confidence interval.

### Monkey experiments – Behavioral outcome decoding procedure

MUA and LFP recorded during the task were aligned on target presentation time and sorted according to monkey’s behavioral response (Correct trials or misses). A regularized linear decoder was used to associate, the neuronal activity estimated on a 200ms *post-target* interval (corresponding to the FEF population responses to the visual target presentation) to the two possible behavioral response of subject. During training, the input to the classifier was a 48 elements by N-trial matrix corresponding to the average neuronal response on each recording channel for the time interval of interest for each of the N training trials. The imposed output of the classifier was the behavioral response of these N training trials. During testing, the output of the classifier was estimated for a 48 element vector corresponding to the average neuronal response on each recording channel for the time interval of interest on a testing trial, new to the classifier. This output can be read as a class output, corresponding to one of the two behavioral responses. Supplementary Fig. 1 presents for MUA and LFP signal based decoder accuracies to classify behavioral outcome (hit or miss). Data were averaged on 19 sessions and compared to 1000 random distribution based on trials labels shuffling for each session. To quantify prefrontal encoding stability across session time, we performed cross-temporal decoding analyses along the entire length of the session. A classifier was trained on a given training set of 200 trials from a specific time in the session. For each individual training set, behavioral response categories (correct or miss) were randomly equalized across trials (100 repetitions). Testing was then performed on the next 50 trials and classification performance was computed on these 50 trials. This procedure was iterated by steps of 1 correct trial at a time, to cover the entire session trial range. The classification performance of each testing set of 50 trials was assigned to the Gaussian mean of the presentation time of these 50 trials relative to the session (σ=length of testing set/6, here σ=8.32). This procedure resulted in a time series representing running classification performance as a function of session time.

### Monkey experiments – Attention (x,y) position decoding procedure

Similarly, to the decoding procedure described in the previous section, a regularized linear decoder was used to associate, on correct trials, the neuronal activity estimated on a 400ms pre-target interval to the expected position of subject’s attention(Astrand et al., 2016; De Sousa et al., 2021; Gaillard et al., 2020a). During training, the input to the classifier was a 48 elements by N-trial matrix corresponding to the average neuronal response on each recording channel for the time interval of interest for each of the N training trials. The imposed output of the classifier was the (x,y) coordinates of the cued landmark for each of these N training trials. During testing, the output of the classifier was estimated for a 48 element vector corresponding to the average neuronal response on each recording channel for the time interval of interest on a testing trial, new to the classifier. This output can be read as a continuous (x,y) estimate of attention location or as a class output, corresponding to one of the four possible visual quadrants (Astrand et al., 2016; Gaillard et al., 2020a).

### Monkey experiments – Attention classification procedure

To quantify prefrontal attention stability across session time, we also performed cross-temporal decoding analyses along the length of the session. A classifier was trained on a given training set of 200 successive correct trials from a specific time in the session. For each individual training set, cued spatial categories (position) were randomly equalized across trials (100 repetitions). Testing was then performed on the next 25 correct trials and classification performance was computed on these 25 trials. This procedure was iterated by steps of 1 correct trial at a time, to cover the entire session trial range. Because of the randomized structure of the task and the variable rate of incorrect trials over session time, the actual duration of each training set varied within and across session (4.550 ±0.547 min). To minimize this temporal sampling variability, and have a robust estimate of mean trial distribution, the classification performance of each testing set of 25 trials was assigned to the Gaussian mean of the presentation time of these 25 trials relative to the session (σ=length of testing set/6, here σ=4.166). This procedure resulted in a time series representing running classification performance as a function of session time.

### Monkey experiments – Signal frequency analyses of classification performance time series

Each independent session classification performance time series was interpolated (cubic spline function). This resulted in a fixed sampling frequency of 1000 ±179 sample/hour for an average session duration of 3.526 hours ± 0.6796. These parameters allow the description of frequencies ranging from 0.29 ± 0.515 to 500 ± 89.97 cycle per hour. The spectral analysis of this time series was performed on detrended data using a Morlet Wavelet transform.

### Monkey experiments – Phase analysis

For each session, phase difference distributions was obtained using a cross wavelet transform analysis (Grinsted et al., 2004). Specifically, cross wavelet transform computes at each frequency and each timestamp the relative phase difference between both time series at the same frequency band. Phases distribution were compared across all sessions and both subjects using non parametric circular Kuiper’s test for equal distributions. Phase comparison were performed between time series constructed from comparable timestamps only, defined as follows. Specifically, behavioral performance and decoded behavioral neuronal time series are both computed over hit and miss trials. They are thus estimated at exactly the same timestamps in the session (condition 1). In contrast, decoded attentional time series for MUA or LFP are assessed on correct trials. These time series are thus estimated at exactly the same timestamps (condition 2). However, the timestamps estimated in condition 1 and condition 2 are different. Indeed, the presence of miss trials of varying frequency of occurrence leads to local time compression or expansion of one timestamp series relative to the other. Because phase estimation is a very sensitive metric, phase analysis could only be performed across time series estimated from the same timestamps.

### Monkey experiments – Performance at optimal phase computation

Average performance at optimal phase was estimated independently computed for each session then averaged. Specifically, for each session, identified oscillatory cycle was independently subdivided into 17 phase bins, and a phase to behavioral performance function was calculated. Optimal phase was defined for each session as the absolute phase of the cycle maximizing subject performances(Busch and VanRullen, 2010; Dugué et al., 2014; Gaillard et al., 2020a). Subsequently, phase to behavioral phases functions were aligned across sessions and averaged to compute averaged relative increase in performance at optimal versus suboptimal phase of the oscillation. Central points (0 phase, by definition maximal) were removed from statistical all analysis.

### Monkey experiments – LFP power and coherency analysis

LFP power spectra were estimated for each channel and each trial using Morlet Wavelet transform over an attentional period ranging from 500 ms post cue presentation to 2000 ms (1.5s epoch). Coherency was computed separately for each hemisphere to avoid possible inter-hemispheric phase differences in LFP frequency content. Coherency was computed using pairwise phase consistency (PPC) performed on a 1 second time epoch (500 ms post cue presentation to 1500 ms) and was based on the fieldtrip *ft_connectivityanalysis* function(Oostenveld et al., 2011).

### Monkey experiments – Spike LFP phase coupling

For each spike within a 1s time window prior to target presentation (on trials with cue to target intervals of 1500ms or higher), we computed the LFP phase estimates at spike time, using pairwise phase consistency estimation (PPC). PPC allows to compute spike LFP phase coupling (SFC) independently of spike and trial number. This was performed using the fieldtrip *ft_spiketriggeredspectrummatlab* function. In order to adapt SFC computation to frequency bands of interest, we used 4, 6 and 8 cycles per frequency to compute frequency phases estimates in respectively the 2-20Hz, 20-50Hz and 50-70Hz frequency bands.

### Monkey experiments – Peak and trough session oscillation estimation

For each session, peak and trough trials were identified based on time series rhythmic variations in session time. First, the behavioral signal was filtered at the specific low-frequency dominating the behavioral performance time series, per session. Then peak (resp. trough) categories were associated to trials occurring on oscillatory peaks (resp. troughs) of the considered time series of the session. All following analysis concerning firing rates, LFP power and coherency and SFC estimates were compared across “peak” and “trough” trials, across all channels and all sessions using non-parametric rank sum tests (n=19 sessions, 2 monkeys).

### *Human experiments* – Human participants

Thirty-five subjects participated in the study (mean age 26.5 ± 8.35 years, fifteen females). Among these, nineteen subjects (age 25.2 ± 2.0, five females) were recorded with a video eye tracker (Eyelink 1000, SR Research) while performing the task. All participants were totally naïve to the purpose of the experiment, reported no history of neurological or psychiatric disorders, had normal or corrected-to-normal vision and participated voluntarily after providing signed informed consent. Inclusion criteria included right-handedness, which was self-reported in all participants. All individual data was anonymized. All human experiments were performed in accordance with relevant guidelines and regulations (the declaration of Helsinski). All experimental protocols were approved by the CNRS (research institution acting as promotor) and the CPP-Sud-Est (acting as licensing committee). The project authorization is #ID RCB 2017-A03612-51.

### *Human experiments* – Behavioral task

Participants were seated in a comfortable chair in a dark room with their head positioned on a chin-rest and their eyes directed towards a computer screen (24 inches), 56 cm away. A custom-made script using Psychotoolbox(Brainard, 1997) in MATLAB (Mathworks) was used to present the stimuli and record performance. Participants were asked to perform a vigilance task. At the beginning of each trial, participants were exposed to a gray-background screen with one-hundred black and white Gabor patches randomly distributed over the screen, of varying orientation, all moving toward different directions at a constant very slow velocity. After a randomized and unpredictable time period that spanned between 5 and 50 seconds, one of these gabor patches (target stimulus) started to shiver. Participants’ task consisted in finding this target gabor stimulus, freely exploring the screen with their eyes. Participants were instructed to point toward the target using a mouse-controlled cursor and produce a left button press, as fast as possible. The subjects’ response initiated a new trial. Participants had to perform consecutive trials during 75 minutes. Importantly, subjects were instructed that there would be no rest periods. For a subset of subjects, during the task, a remote gaze tracking capture system (Eyelink 1000, SR Research) was used to continuously record eye movements, blinking and pupil size. These data were recorded at a sampling rate of 500 Hz. The eye tracker was calibrated for each subject before the main task was initiated.

### Human experiments – Behavioral performance as a function of session time and spectral analyses

Participants’ performance was measured by studying the fluctuations of the target detection time (DT) along the task. Detection times are defined as the time between the onset of the target stimulus and the subject’s button press. In this way, we obtained the time series of the detection times. In order to study the spectral decomposition of this time series, we first equalized its sampling rate to 1000 samples per hour using a least-square interpolation method, allowing a good time resolution for frequency analysis. Spectral analyses were the same as those performed on the monkey behavioral data. Specifically, the spectral analysis of this time series was performed on detrended data using a Morlet Wavelet transform. Standard error represented on the power spectrum is calculated along the time dimension of the wavelet analysis. We established statistical significance by reshuffling trials presentation time across the session (1000 times) and generating a random distribution of signal spectral content. Significance was then computed by comparing each session power spectrum to its corresponding 95% confidence interval.

### *Human experiments* – Pupil size *as a function of session time and spectral analyses*

Pupil size was recorded using the video eye tracking system. Blinking-related artifact was removed as follows. A cubic spline interpolation method was applied to fill in data between blink onset and blink offset (defined as a sharp increase and decrease in pupil size). After removing these blinking artifacts, pupil size time series was down-sampled to 1 Hz and data was then processed using the same method as the behavioral detection time series. Namely, the spectral analysis of this time series was performed on detrended data using a Morlet Wavelet transform. Standard error represented on the power spectrum is calculated along the time dimension of the wavelet analysis. We established statistical significance by reshuffling trials presentation time across the session (1000 times) and generating a random distribution of signal spectral content. Significance was then computed by comparing each session power spectrum to its corresponding 95% confidence interval.

Pupil Dilation Reflects the Creation and Retrieval of Memories – Stephen D. Goldinger, Megan H. Papesh, 2012.

## Supplementary figures

**Figure S1:**
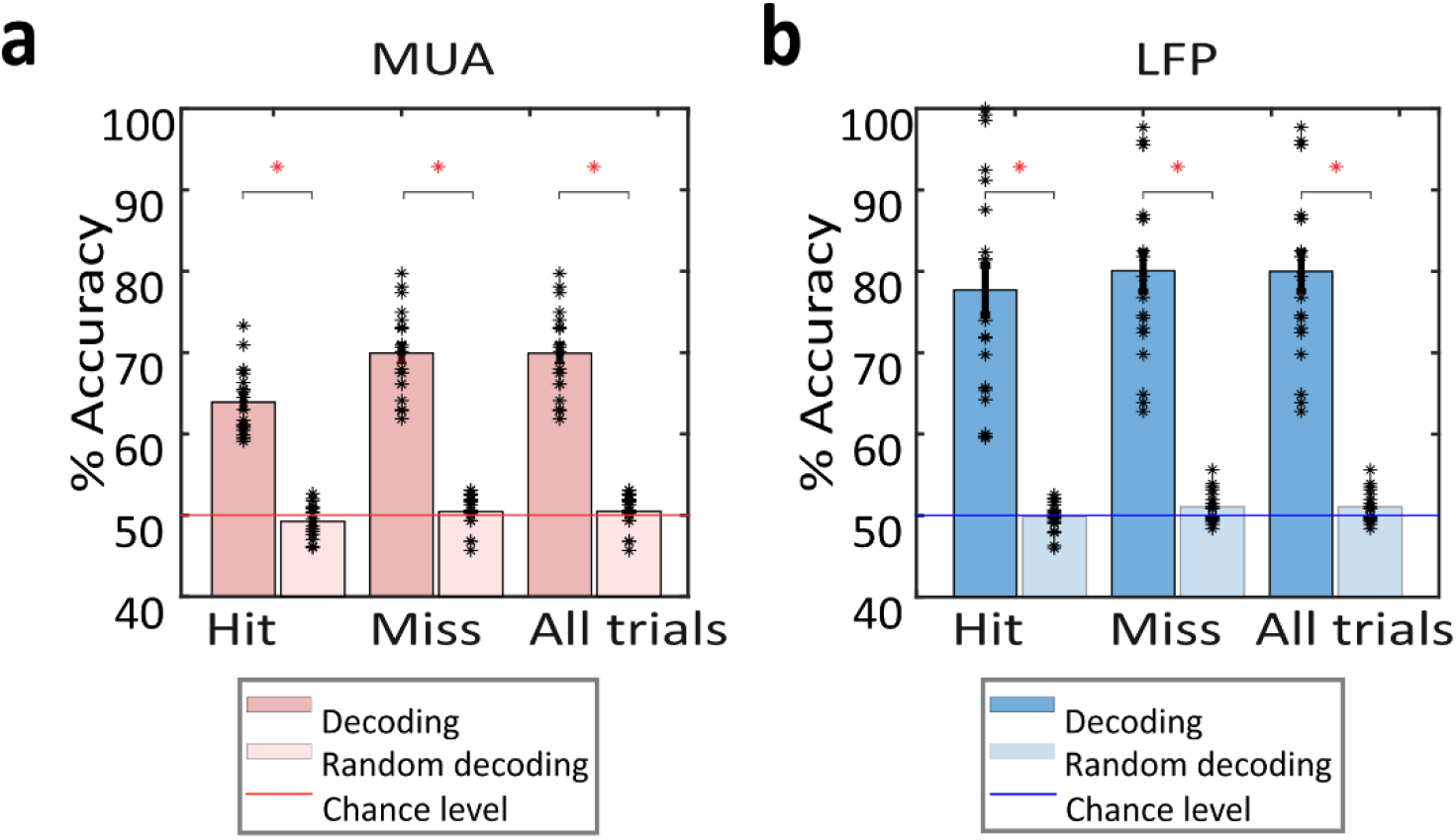
Hit vs. Miss decoding accuracies from prefrontal MUA and LFP signals. (A) Decoding accuracies for Hit, Miss or all trials (Hit and Miss) classification from prefrontal MUA presented in pink compare to a random decoding presented in white (n=19; ranksum test – *p value < 0.0001). (B) Decoding accuracies for Hit, Miss or both trials (Hit and Miss) classification from prefrontal LFP frequency content presented in blue compare to a random decoding presented in white (n=19; ranksum test – *p value < 0.0001).

**Figure S2:**
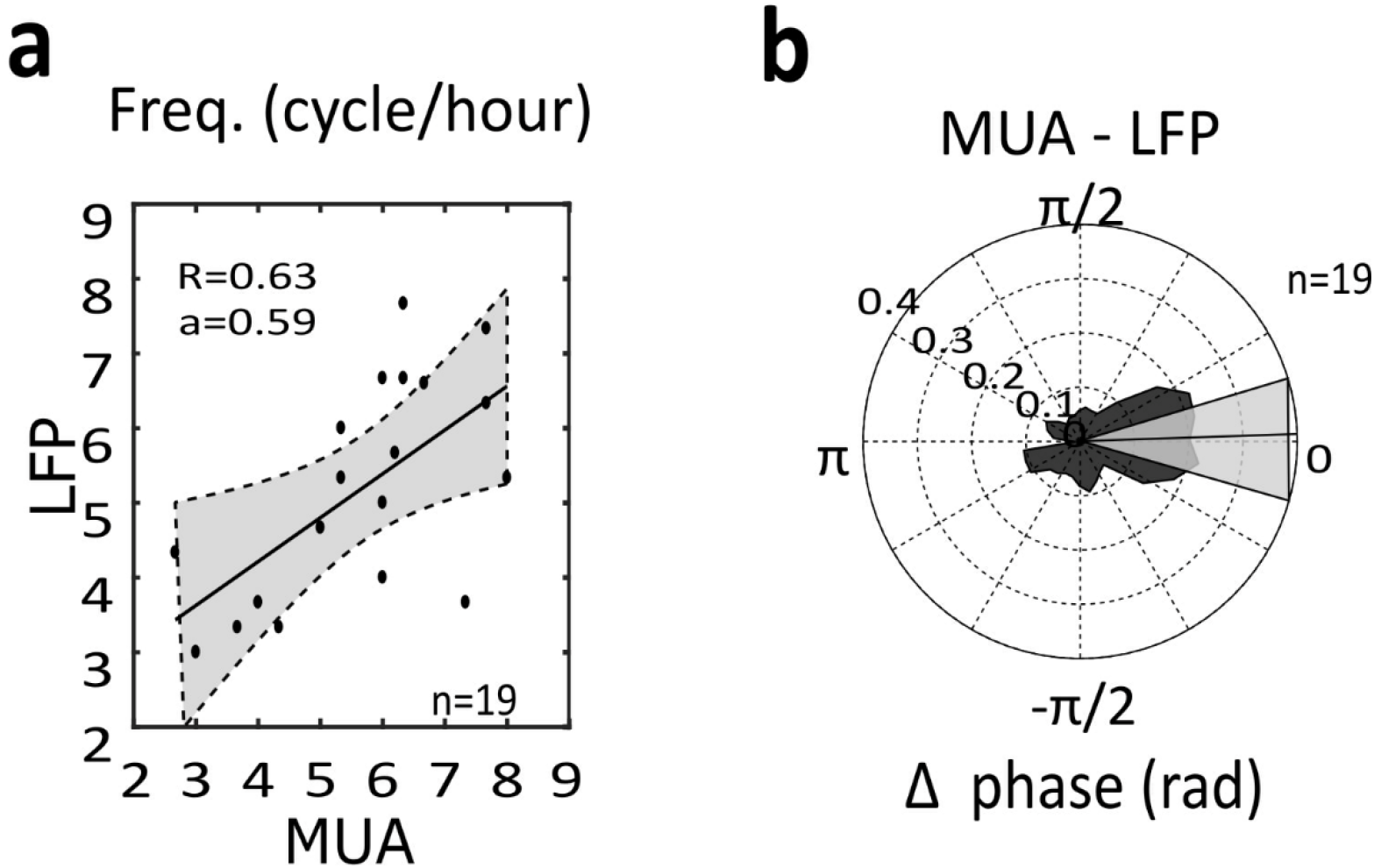
Ultra-slow rhythmic fluctuations in accuracies at target decoding on hit vs. miss trials from prefrontal MUA and LFP signals. (A) Correlation between ultra-slow oscillations identified in LFP and MUA hit vs. miss decoding accuracies (Spearman correlation, p<0.01) and corresponding phase difference (B) (circular Kuiper’s test p<0.05).

**Figure S3:**
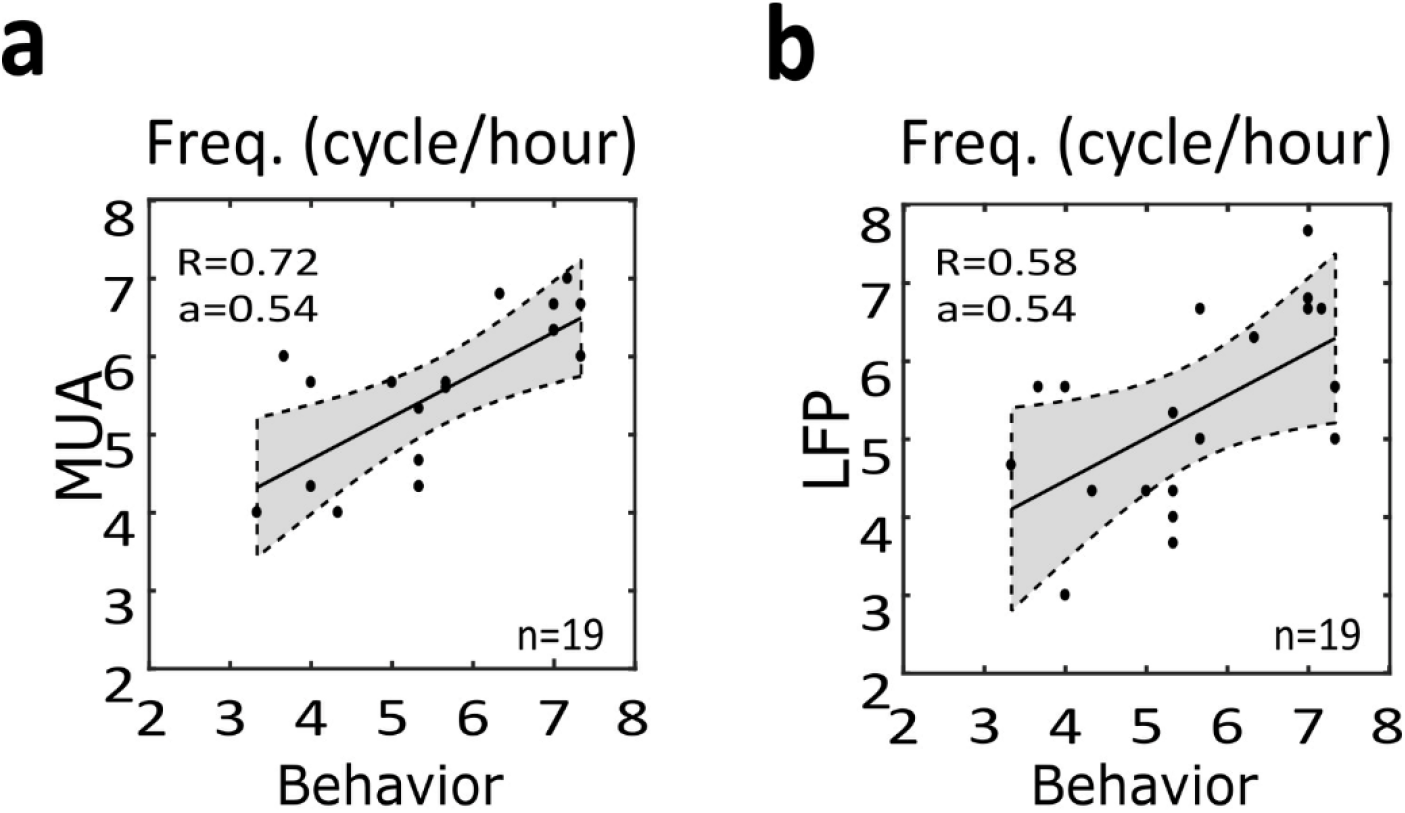
Ultra-slow rhythmic fluctuations in accuracies at decoding attention-related information from prefrontal MUA and LFP signals. (A) Correlation between ultra-slow oscillations identified in MUA spatial decoding accuracies and in behavioral performance (Spearman correlation, p<0.01) and corresponding phase difference (B) (circular Kuiper’s test p<0.05).

**Figure S4:**
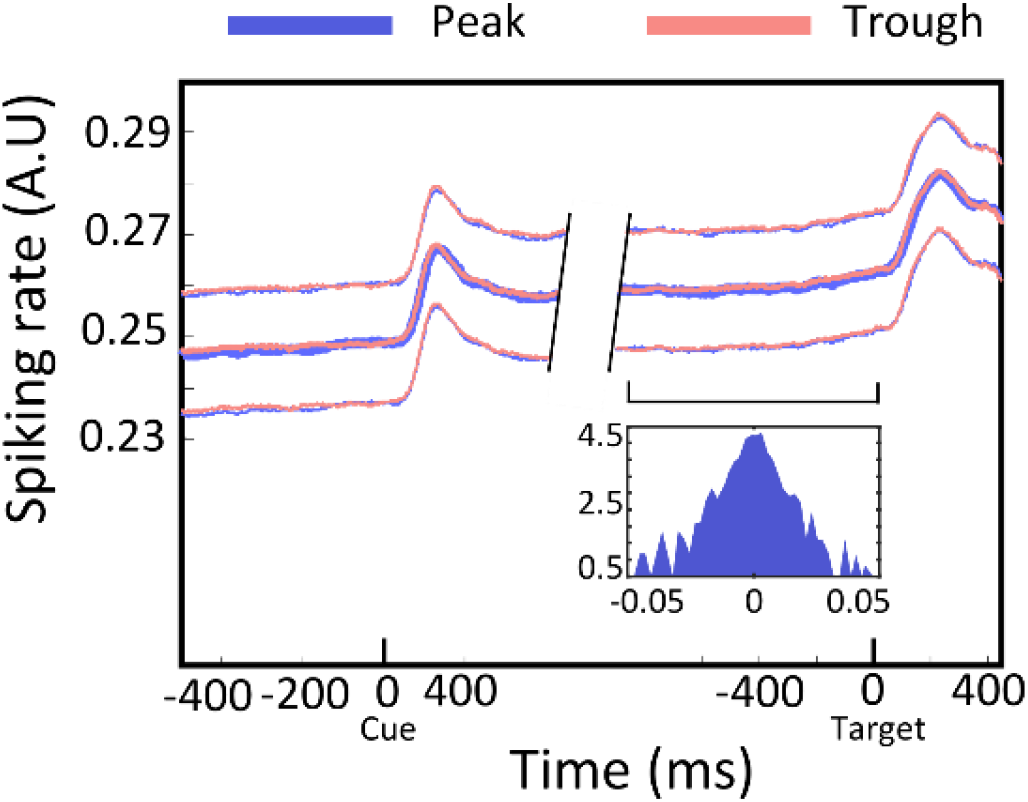
Ultra-slow rhythm does not impact *averaged FEF MUA response to Cue and Target presentation.* Averaged MUA aligned on Cue and Target presentation for trials belonging to either optimal phase of the oscillation (Peak, Blue, mean +-SE) or sub-optimal phase of the oscillation (Peak, pink, mean +-SE), Wilcoxon rank sum test p>0.5. Inset represents Modulation Index for MUA responses corresponding to all channels (n=912) analyzed (see method).

## References

Abrahamyan, A., Silva, L.L., Dakin, S.C., Carandini, M., and Gardner, J.L. (2016). Adaptable history biases in human perceptual decisions. PNAS 113, E3548–E3557.

Allen, W.E., Chen, M.Z., Pichamoorthy, N., Tien, R.H., Pachitariu, M., Luo, L., and Deisseroth, K. (2019). Thirst regulates motivated behavior through modulation of brainwide neural population dynamics. Science 364.

Arieli, A., Sterkin, A., Grinvald, A., and Aertsen, A. (1996). Dynamics of ongoing activity: explanation of the large variability in evoked cortical responses. Science 273, 1868–1871.

Aston-Jones, G., and Cohen, J.D. (2005). An integrative theory of locus coeruleus-norepinephrine function: adaptive gain and optimal performance. Annu Rev Neurosci 28, 403–450.

Astrand, E., Wardak, C., Baraduc, P., and Ben Hamed, S. (2016). Direct Two-Dimensional Access to the Spatial Location of Covert Attention in Macaque Prefrontal Cortex. Curr. Biol. 26, 1699–1704.

Astrand, E., Wardak, C., and Ben Hamed, S. (2020). Neuronal population correlates of target selection and distractor filtering. NeuroImage 209, 116517.

B, L., S, S., and G, G. (2012). Pupillometry: A Window to the Preconscious? Perspectives on Psychological Science: A Journal of the Association for Psychological Science 7.

Bagherzadeh, Y., Baldauf, D., Pantazis, D., and Desimone, R. (2020). Alpha Synchrony and the Neurofeedback Control of Spatial Attention. Neuron 105, 577–587.e5.

Barttfeld, P., Uhrig, L., Sitt, J.D., Sigman, M., Jarraya, B., and Dehaene, S. (2015). Signature of consciousness in the dynamics of resting-state brain activity. Proc Natl Acad Sci U S A 112, 887–892.

Beaman, C.B., Eagleman, S.L., and Dragoi, V. (2017). Sensory coding accuracy and perceptual performance are improved during the desynchronized cortical state. Nature Communications 8, 1308.

Beane, M., and Marrocco, R.T. (2004). Norepinephrine and acetylcholine mediation of the components of reflexive attention: implications for attention deficit disorders. Prog Neurobiol 74, 167–181.

Beatty, J., and Lucero-Wagoner, B. (2000). The pupillary system. In Handbook of Psychophysiology, 2nd Ed, (New York, NY, US: Cambridge University Press), pp. 142–162.

Bello, F.D., Hassen, S.B.H., Astrand, E., and Hamed, S.B. (2020). Selection and suppression of visual information in the macaque prefrontal cortex. BioRxiv 2020.03.25.007922.

Bollimunta, A., Chen, Y., Schroeder, C.E., and Ding, M. (2008). Neuronal Mechanisms of Cortical Alpha Oscillations in Awake-Behaving Macaques. J. Neurosci. 28, 9976–9988.

Bonnefond, A., Doignon-Camus, N., Touzalin-Chretien, P., and Dufour, A. (2010). Vigilance and intrinsic maintenance of alert state: An ERP study. Behavioural Brain Research 211, 185–190.

Brainard, D.H. (1997). The Psychophysics Toolbox. Spat Vis 10, 433–436.

Busch, N.A., and VanRullen, R. (2010). Spontaneous EEG oscillations reveal periodic sampling of visual attention. Proc. Natl. Acad. Sci. U.S.A. 107, 16048–16053.

Busch, N.A., Dubois, J., and VanRullen, R. (2009). The phase of ongoing EEG oscillations predicts visual perception. J. Neurosci. 29, 7869–7876.

Buschman, T.J., and Miller, E.K. (2007a). Top-down versus bottom-up control of attention in the prefrontal and posterior parietal cortices. Science 315, 1860–1862.

Buschman, T.J., and Miller, E.K. (2007b). Top-down versus bottom-up control of attention in the prefrontal and posterior parietal cortices. Science 315, 1860–1862.

Buschman, T.J., and Miller, E.K. (2009). Serial, Covert, Shifts of Attention during Visual Search are Reflected by the Frontal Eye Fields and Correlated with Population Oscillations. Neuron 63, 386–396.

Chalk, M., Herrero, J.L., Gieselmann, M.A., Delicato, L.S., Gotthardt, S., and Thiele, A. (2010). Attention Reduces Stimulus-Driven Gamma Frequency Oscillations and Spike Field Coherence in V1. Neuron 66, 114–125.

Chan, R.W., Leong, A.T.L., Ho, L.C., Gao, P.P., Wong, E.C., Dong, C.M., Wang, X., He, J., Chan, Y.-S., Lim, L.W., et al. (2017). Low-frequency hippocampal-cortical activity drives brain-wide resting-state functional MRI connectivity. Proc Natl Acad Sci U S A 114, E6972–E6981.

Compton, R.J., Gearinger, D., and Wild, H. (2019). The wandering mind oscillates: EEG alpha power is enhanced during moments of mind-wandering. Cogn Affect Behav Neurosci 19, 1184–1191.

Cowley, B.R., Snyder, A.C., Acar, K., Williamson, R.C., Yu, B.M., and Smith, M.A. (2020). Slow Drift of Neural Activity as a Signature of Impulsivity in Macaque Visual and Prefrontal Cortex. Neuron 108, 551–567.e8.

De Sousa, C., Gaillard, C., Di Bello, C., Ben Hadj Hassen, F., and Ben Hamed, S. (2021). Behavioral validation of novel high resolution attention decoding method from multi-units & local field potentials. NeuroImage 231, 117853.

Demertzi, A., Tagliazucchi, E., Dehaene, S., Deco, G., Barttfeld, P., Raimondo, F., Martial, C., Fernández-Espejo, D., Rohaut, B., Voss, H.U., et al. (2019). Human consciousness is supported by dynamic complex patterns of brain signal coordination. Science Advances 5, eaat7603.

van Dijk, H., Schoffelen, J.-M., Oostenveld, R., and Jensen, O. (2008). Prestimulus oscillatory activity in the alpha band predicts visual discrimination ability. J Neurosci 28, 1816–1823.

Dragone, A., Lasaponara, S., Pinto, M., Rotondaro, F., De Luca, M., and Doricchi, F. (2018). Expectancy modulates pupil size during endogenous orienting of spatial attention. Cortex 102, 57–66.

Dugué, L., and VanRullen, R. (2017). Transcranial Magnetic Stimulation Reveals Intrinsic Perceptual and Attentional Rhythms. Front Neurosci 11.

Dugué, L., Marque, P., and VanRullen, R. (2014). Theta Oscillations Modulate Attentional Search Performance Periodically. Journal of Cognitive Neuroscience 27, 1–14.

Dugué, L., Roberts, M., and Carrasco, M. (2016). Attention Reorients Periodically. Curr Biol 26, 1595–1601.

Ebitz, R.B., and Moore, T. (2017). Selective Modulation of the Pupil Light Reflex by Microstimulation of Prefrontal Cortex. J. Neurosci. 37, 5008–5018.

Ekstrom, L.B., Roelfsema, P.R., Arsenault, J.T., Bonmassar, G., and Vanduffel, W. (2008). Bottom-up dependent gating of frontal signals in early visual cortex. Science 321, 414–417.

Eldar, E., Cohen, J.D., and Niv, Y. (2013). The effects of neural gain on attention and learning. Nat Neurosci 16, 1146–1153.

Engel, T.A., Steinmetz, N.A., Gieselmann, M.A., Thiele, A., Moore, T., and Boahen, K. (2016). Selective modulation of cortical state during spatial attention. Science 354, 1140–1144.

Everitt, B.J., and Robbins, T.W. (1997). Central cholinergic systems and cognition. Annu Rev Psychol 48, 649–684.

Fiebelkorn, I.C., Pinsk, M.A., and Kastner, S. (2018a). A Dynamic Interplay within the Frontoparietal Network Underlies Rhythmic Spatial Attention. Neuron 99, 842–853.e8.

Fiebelkorn, I.C., Pinsk, M.A., and Kastner, S. (2018b). A Dynamic Interplay within the Frontoparietal Network Underlies Rhythmic Spatial Attention. Neuron 99, 842–853.e8.

Fiebelkorn, I.C., Pinsk, M.A., and Kastner, S. (2019). The mediodorsal pulvinar coordinates the macaque fronto-parietal network during rhythmic spatial attention. Nature Communications 10, 215.

Fries, P. (2005). A mechanism for cognitive dynamics: neuronal communication through neuronal coherence. Trends in Cognitive Sciences 9, 474–480.

Fries, P. (2015a). Rhythms For Cognition: Communication Through Coherence. Neuron 88, 220–235.

Fries, P. (2015b). Rhythms For Cognition: Communication Through Coherence. Neuron 88, 220–235.

Fries, P., Reynolds, J.H., Rorie, A.E., and Desimone, R. (2001). Modulation of oscillatory neuronal synchronization by selective visual attention. Science 291, 1560–1563.

Fries, P., Womelsdorf, T., Oostenveld, R., and Desimone, R. (2008). The Effects of Visual Stimulation and Selective Visual Attention on Rhythmic Neuronal Synchronization in Macaque Area V4. J Neurosci 28, 4823–4835.

Gaillard, C., Ben Hadj Hassen, S., Di Bello, F., Bihan-Poudec, Y., VanRullen, R., and Ben Hamed, S. (2020a). Prefrontal attentional saccades explore space rhythmically. Nature Communications 11, 925.

Gaillard, C., Ben Hadj Hassen, S., Di Bello, F., Bihan-Poudec, Y., VanRullen, R., and Ben Hamed, S. (2020b). Prefrontal attentional saccades explore space rhythmically. Nat Commun 11, 925.

Gregoriou, G.G., Gotts, S.J., Zhou, H., and Desimone, R. (2009a). High-frequency, long-range coupling between prefrontal and visual cortex during attention. Science 324, 1207–1210.

Gregoriou, G.G., Gotts, S.J., Zhou, H., and Desimone, R. (2009b). High-frequency, long-range coupling between prefrontal and visual cortex during attention. Science 324, 1207–1210.

Grinsted, A., Moore, J.C., and Jevrejeva, S. (2004). Application of the cross wavelet transform and wavelet coherence to geophysical time series. Nonlinear Processes in Geophysics 11, 561–566.

Guo, Z., Chen, R., Zhang, K., Pan, Y., and Wu, J. (2016). The Impairing Effect of Mental Fatigue on Visual Sustained Attention under Monotonous Multi-Object Visual Attention Task in Long Durations: An Event-Related Potential Based Study. PLOS ONE 11, e0163360.

Gutnisky, D.A., Beaman, C., Lew, S.E., and Dragoi, V. (2017). Cortical response states for enhanced sensory discrimination. Elife 6.

Hassen, S.B.H., Wardak, C., and Hamed, S.B. (2019). Rhythmic variations in prefrontal interneuronal correlations, their underlying mechanisms and their behavioral correlates. BioRxiv 784850.

Horschig, J.M., Oosterheert, W., Oostenveld, R., and Jensen, O. (2015). Modulation of Posterior Alpha Activity by Spatial Attention Allows for Controlling A Continuous Brain-Computer Interface. Brain Topogr 28, 852–864.

Huang, N., and Elhilali, M. Push-pull competition between bottom-up and top-down auditory attention to natural soundscapes. ELife 9.

Huang, Z., Zhang, J., Wu, J., Mashour, G.A., and Hudetz, A.G. (2020). Temporal circuit of macroscale dynamic brain activity supports human consciousness. Sci Adv 6, eaaz0087.

van der Hulst, M. (2003). Long workhours and health. Scand J Work Environ Health 29, 171–188.

Ibos, G., Duhamel, J.-R., and Hamed, S.B. (2013). A Functional Hierarchy within the Parietofrontal Network in Stimulus Selection and Attention Control. J. Neurosci. 33, 8359–8369.

Jasper, A.I., Tanabe, S., and Kohn, A. (2019). Predicting Perceptual Decisions Using Visual Cortical Population Responses and Choice History. J. Neurosci. 39, 6714–6727.

Jia, X., and Kohn, A. (2011). Gamma Rhythms in the Brain. PLoS Biol 9.

Johnson, J.V., and Lipscomb, J. (2006). Long working hours, occupational health and the changing nature of work organization. Am J Ind Med 49, 921–929.

Joshi, S., Li, Y., Kalwani, R.M., and Gold, J.I. (2016). Relationships between Pupil Diameter and Neuronal Activity in the Locus Coeruleus, Colliculi, and Cingulate Cortex. Neuron 89, 221–234.

Just, M.A., Carpenter, P.A., and Miyake, A. (2003). Neuroindices of cognitive workload: Neuroimaging, pupillometric and event-related potential studies of brain work. Theoretical Issues in Ergonomics Science 4, 56–88.

Kembro, J.M., Lihoreau, M., Garriga, J., Raposo, E.P., and Bartumeus, F. (2019). Bumblebees learn foraging routes through exploitation-exploration cycles. J R Soc Interface 16, 20190103.

Kim, S.-Y., and Lim, W. (2013). Sparsely-synchronized brain rhythm in a small-world neural network. Journal of the Korean Physical Society 63, 104–113.

Klein, R., and Armitage, R. (1979). Rhythms in human performance: 1 1/2-hour oscillations in cognitive style. Science 204, 1326–1328.

Knyazev, G.G., Slobodskoj-Plusnin, J.Y., Bocharov, A.V., and Pylkova, L.V. (2011). The default mode network and EEG α oscillations: an independent component analysis. Brain Res 1402, 67–79.

Kramer, A. (1990). Physiological metrics of mental workload: A review of recent progress.

Leopold, D.A., and Logothetis, N.K. (1998). Microsaccades differentially modulate neural activity in the striate and extrastriate visual cortex. Exp Brain Res 123, 341–345.

Li, J.M., Bentley, W.J., Snyder, A.Z., Raichle, M.E., and Snyder, L.H. (2015). Functional connectivity arises from a slow rhythmic mechanism. Proc Natl Acad Sci U S A 112, E2527–2535.

Liu, Y., and Tanaka, H. (2002). Overtime work, insufficient sleep, and risk of non-fatal acute myocardial infarction in Japanese men. Occup Environ Med 59, 447–451.

Liu, Y., Rodenkirch, C., Moskowitz, N., Schriver, B., and Wang, Q. (2017). Dynamic Lateralization of Pupil Dilation Evoked by Locus Coeruleus Activation Results from Sympathetic, Not Parasympathetic, Contributions. Cell Rep 20, 3099–3112.

Lockley, S., Cronin, J., Flynn-Evans, E., Cade, B., Lee, C., Landrigan, C., Rothschild, J., Katz, J., Lilly, C., Stone, P., et al. (2004). Effect of Reducing Interns’ Weekly Work Hours on Sleep and Attentional Failures. The New England Journal of Medicine 351, 1829–1837.

Marcora, S.M., Staiano, W., and Manning, V. (2009). Mental fatigue impairs physical performance in humans. J Appl Physiol (1985) 106, 857–864.

McCormick, D.A. (1992). Neurotransmitter actions in the thalamus and cerebral cortex and their role in neuromodulation of thalamocortical activity. Progress in Neurobiology 39, 337–388.

McGinley, M.J., Vinck, M., Reimer, J., Batista-Brito, R., Zagha, E., Cadwell, C.R., Tolias, A.S., Cardin, J.A., and McCormick, D.A. (2015). Waking State: Rapid Variations Modulate Neural and Behavioral Responses. Neuron 87, 1143–1161.

Mi, Y., Liao, X., Huang, X., Zhang, L., Gu, W., Hu, G., and Wu, S. (2013). Long-period rhythmic synchronous firing in a scale-free network. Proc Natl Acad Sci U S A 110, E4931–4936.

Milton, R., Shahidi, N., and Dragoi, V. (2020). Dynamic states of population activity in prefrontal cortical networks of freely-moving macaque. Nature Communications 11, 1948.

Mo, J., Schroeder, C.E., and Ding, M. (2011). Attentional Modulation of Alpha Oscillations in Macaque Inferotemporal Cortex. J Neurosci 31, 878–882.

Mo, J., Liu, Y., Huang, H., and Ding, M. (2013). Coupling between visual alpha oscillations and default mode activity. Neuroimage 68, 112–118.

Moore, T., and Zirnsak, M. (2017). Neural Mechanisms of Selective Visual Attention. Annu Rev Psychol 68, 47–72.

Nassar, M.R., Rumsey, K.M., Wilson, R.C., Parikh, K., Heasly, B., and Gold, J.I. (2012). Rational regulation of learning dynamics by pupil-linked arousal systems. Nat Neurosci 15, 1040–1046.

Oby, E.R., Golub, M.D., Hennig, J.A., Degenhart, A.D., Tyler-Kabara, E.C., Yu, B.M., Chase, S.M., and Batista, A.P. (2019). New neural activity patterns emerge with long-term learning. Proc Natl Acad Sci U S A 116, 15210–15215.

Okun, M., Steinmetz, N.A., Cossell, L., Iacaruso, M.F., Ko, H., Barthó, P., Moore, T., Hofer, S.B., Mrsic-Flogel, T.D., Carandini, M., et al. (2015). Diverse coupling of neurons to populations in sensory cortex. Nature 521, 511–515.

Okun, M., Steinmetz, N.A., Lak, A., Dervinis, M., and Harris, K.D. (2019). Distinct Structure of Cortical Population Activity on Fast and Infraslow Timescales. Cerebral Cortex 29, 2196–2210.

Oostenveld, R., Fries, P., Maris, E., and Schoffelen, J.-M. (2011). FieldTrip: Open source software for advanced analysis of MEG, EEG, and invasive electrophysiological data. Comput Intell Neurosci 2011, 156869.

Palva, J.M., and Palva, S. (2011). Roles of multiscale brain activity fluctuations in shaping the variability and dynamics of psychophysical performance. Prog Brain Res 193, 335–350.

Paneri, S., and Gregoriou, G.G. (2017). Top-Down Control of Visual Attention by the Prefrontal Cortex. Functional Specialization and Long-Range Interactions. Front Neurosci 11.

Parto Dezfouli, M., Khamechian, M.B., Treue, S., Esghaei, M., and Daliri, M.R. (2018). Neural Activity Predicts Reaction in Primates Long Before a Behavioral Response. Front Behav Neurosci 12.

Pisupati, S., Chartarifsky-Lynn, L., Khanal, A., and Churchland, A.K. Lapses in perceptual decisions reflect exploration. ELife 10.

Posner, M.I. (2016). Orienting of attention: Then and now. Q J Exp Psychol (Hove) 69, 1864–1875.

Proctor, S.P., White, R.F., Robins, T.G., Echeverria, D., and Rocskay, A.Z. (1996). Effect of overtime work on cognitive function in automotive workers. Scand J Work Environ Health 22, 124–132.

Rabinowitz, N.C., Goris, R.L., Cohen, M., and Simoncelli, E.P. (2015). Attention stabilizes the shared gain of V4 populations. ELife 4, e08998.

Reimer, J., Froudarakis, E., Cadwell, C.R., Yatsenko, D., Denfield, G.H., and Tolias, A.S. (2014). Pupil fluctuations track fast switching of cortical states during quiet wakefulness. Neuron 84, 355–362.

Reynaud, A.J., Froesel, M., Guedj, C., Ben Hadj Hassen, S., Cléry, J., Meunier, M., Ben Hamed, S., and Hadj-Bouziane, F. (2019). Atomoxetine improves attentional orienting in a predictive context. Neuropharmacology 150, 59–69.

Ros, T., Michela, A., Bellman, A., Vuadens, P., Saj, A., and Vuilleumier, P. (2017). Increased Alpha-Rhythm Dynamic Range Promotes Recovery from Visuospatial Neglect: A Neurofeedback Study. Neural Plast 2017.

Saj, A., Ros, T., Michela, A., and Vuilleumier, P. (2018). Effect of a single early EEG neurofeedback training on remediation of spatial neglect in the acute phase. Ann Phys Rehabil Med 61, 111–112.

Sekine, M., Chandola, T., Martikainen, P., McGeoghegan, D., Marmot, M., and Kagamimori, S. (2006). Explaining social inequalities in health by sleep: the Japanese civil servants study. Journal of Public Health 28, 63–70.

Shields, M. (1999). Long working hours and health. Health Rep 11, 33–48(Eng); 37-55(Fre).

Siegel, M., Donner, T.H., Oostenveld, R., Fries, P., and Engel, A.K. (2008). Neuronal synchronization along the dorsal visual pathway reflects the focus of spatial attention. Neuron 60, 709–719.

Sirois, S., and Brisson, J. (2014). Pupillometry. WIREs Cognitive Science 5, 679–692.

Smallwood, J., and Schooler, J.W. (2015). The science of mind wandering: empirically navigating the stream of consciousness. Annu Rev Psychol 66, 487–518.

Snyder, A.C., Morais, M.J., and Smith, M.A. (2016). Dynamics of excitatory and inhibitory networks are differentially altered by selective attention. J Neurophysiol 116, 1807–1820.

Sokejima, S., and Kagamimori, S. (1998). Working hours as a risk factor for acute myocardial infarction in Japan: case-control study. BMJ 317, 775–780.

Steinmetz, N.A., and Moore, T. (2014). Eye movement preparation modulates neuronal responses in area V4 when dissociated from attentional demands. Neuron 83, 496–506.

Stringer, C., Pachitariu, M., Steinmetz, N., Carandini, M., and Harris, K.D. (2019). High-dimensional geometry of population responses in visual cortex. Nature 571, 361–365.

Thomson, D.R., Besner, D., and Smilek, D. (2015). A resource-control account of sustained attention: evidence from mind-wandering and vigilance paradigms. Perspect Psychol Sci 10, 82–96.

Tremblay, S., Doucet, G., Pieper, F., Sachs, A., and Martinez-Trujillo, J. (2015). Single-Trial Decoding of Visual Attention from Local Field Potentials in the Primate Lateral Prefrontal Cortex Is Frequency-Dependent. J. Neurosci. 35, 9038–9049.

Urai, A.E., de Gee, J.W., Tsetsos, K., and Donner, T.H. (2019). Choice history biases subsequent evidence accumulation. ELife 8, e46331.

VanRullen, R. (2018). Attention Cycles. Neuron 99, 632–634.

Vernet, M., Stengel, C., Quentin, R., Amengual, J.L., and Valero-Cabré, A. (2019). Entrainment of local synchrony reveals a causal role for high-beta right frontal oscillations in human visual consciousness. Sci Rep 9, 14510.

Vinck, M., van Wingerden, M., Womelsdorf, T., Fries, P., and Pennartz, C.M.A. (2010). The pairwise phase consistency: A bias-free measure of rhythmic neuronal synchronization. NeuroImage 51, 112–122.

Vinck, M., Batista-Brito, R., Knoblich, U., and Cardin, J.A. (2015). Arousal and locomotion make distinct contributions to cortical activity patterns and visual encoding. Neuron 86, 740–754.

Virtanen, M., Singh-Manoux, A., Ferrie, J.E., Gimeno, D., Marmot, M.G., Elovainio, M., Jokela, M., Vahtera, J., and Kivimäki, M. (2009). Long Working Hours and Cognitive Function. Am J Epidemiol 169, 596–605.

van Vugt, B., Dagnino, B., Vartak, D., Safaai, H., Panzeri, S., Dehaene, S., and Roelfsema, P.R. (2018). The threshold for conscious report: Signal loss and response bias in visual and frontal cortex. Science 360, 537–542.

Warm, J.S., Parasuraman, R., and Matthews, G. (2008). Vigilance Requires Hard Mental Work and Is Stressful. Hum Factors 50, 433–441.

Wasmuht, D.F., Parker, A.J., and Krug, K. (2019). Interneuronal correlations at longer time scales predict decision signals for bistable structure-from-motion perception. Scientific Reports 9, 11449.

Weissman, D.H., Roberts, K.C., Visscher, K.M., and Woldorff, M.G. (2006). The neural bases of momentary lapses in attention. Nat Neurosci 9, 971–978.

van der Wel, P., and van Steenbergen, H. (2018). Pupil dilation as an index of effort in cognitive control tasks: A review. Psychon Bull Rev 25, 2005–2015.

Yüzgeç, Ö., Prsa, M., Zimmermann, R., and Huber, D. (2018). Pupil Size Coupling to Cortical States Protects the Stability of Deep Sleep via Parasympathetic Modulation. Curr Biol 28, 392–400.e3.

Zareian, B., Daliri, M.R., Maboudi, K., Moghaddam, H.A., Treue, S., and Esghaei, M. (2018). Attention enhances LFP phase coherence in macaque visual cortex, improving sensory processing. BioRxiv 499756.

Zareian, B., Maboudi, K., Daliri, M.R., Abrishami Moghaddam, H., Treue, S., and Esghaei, M. (2020). Attention strengthens across-trial pre-stimulus phase coherence in visual cortex, enhancing stimulus processing. Sci Rep 10.

